# Inferring Patterns of Hybridization and Polyploidy in the Plant Genus *Penstemon* (Plantaginaceae)

**DOI:** 10.1101/2020.09.04.283093

**Authors:** Paul D. Blischak, Coleen E. Thompson, Emiko M. Waight, Laura S. Kubatko, Andrea D. Wolfe

## Abstract

Reticulate evolutionary events are hallmarks of plant phylogeny, and are increasingly recognized as common occurrences in other branches of the Tree of Life. However, inferring the evolutionary history of admixed lineages presents a difficult challenge for systematists due to genealogical discordance caused by both incomplete lineage sorting (ILS) and hybridization. Methods that accommodate both of these processes are continuing to be developed, but they often do not scale well to larger numbers of species. An additional complicating factor for many plant species is the occurrence of whole genome duplication (WGD), which can have various outcomes on the genealogical history of haplotypes sampled from the genome. In this study, we sought to investigate patterns of hybridization and WGD in two subsections from the genus *Penstemon* (Plantaginaceae; subsect. *Humiles* and *Proceri*), a speciose group of angiosperms that has rapidly radiated across North America. Species in subsect. *Humiles* and *Proceri* occur primarily in the Pacific Northwest of the United States, occupying habitats such as mesic, subalpine meadows, as well as more well-drained substrates at varying elevations. Ploidy levels in the subsections range from diploid to hexaploid, and it is hypothesized that most of the polyploids are hybrids (i.e., allopolyploids). To estimate phylogeny in these groups, we first developed a method for estimating quartet concordance factors (QCFs) from multiple sequences sampled per lineage, allowing us to model all haplotypes from a polyploid. QCFs represent the proportion of gene trees that support a particular species quartet relationship, and are used for species network estimation in the program SNaQ (Solís-Lemus & Ané. 2016. *PLoS Genet.* 12:e1005896). Using phased haplotypes for nuclear amplicons, we inferred species trees and networks for 38 taxa from *P*. subsect. *Humiles* and *Proceri*. Our phylogenetic analyses recovered two clades comprising a mix of taxa from both subsections, indicating that the current taxonomy for these groups is inconsistent with our estimates of phylogeny. In addition, there was little support for hypotheses regarding the formation of putative allopolyploid lineages. Overall, we found evidence for the effects of both ILS and admixture on the evolutionary history of these species, but were able to evaluate our taxonomic hypotheses despite high levels of gene tree discordance. Our method for estimating QCFs from multiple haplotypes also allowed us to include species of varying ploidy levels in our analyses, which we anticipate will help to facilitate estimation of species networks in other plant groups as well.

## 1 Introduction

Phylogenetic inference with multiple gene sequences has emerged as a dominant paradigm in systematics, with multilocus datasets ranging in size from just a handful of genes, to thousands of loci pulled from whole genomes. Discordant signals from these different gene regions can often be present, however, raising the issue of how to model the incongruence among the sampled gene trees from the underlying species tree. The multispecies coalescent (MSC) model is one approach for species tree estimation from multilocus data that can accommodate gene tree discordance caused by incomplete lineage sorting (ILS) (reviewed in Degnan and Rosenberg 2009). The appeal of the MSC model stems from its connection with concepts in population genetics (Wright-Fisher model; Kingman 1982), and its explicit predictions regarding the amount of gene tree discordance that should be present for a given species tree (Tavaré 1984; Pamilo and Nei 1988; Takahata 1989). Nevertheless, despite the popularity of the coalescent model, it has been shown that it can be a poor fit to empirical data sets (Reid et al. 2014; Gruenstaeudl et al. 2015). A potential reason for the poor performance of the MSC in empirical data is that it only models ILS, leaving other processes that generate genealogical discordance, such as gene flow and hybridization, unaccounted for (Maddison 1997).

An alternative to using the coalescent to model gene tree discordance is to use the concept of a “gene-to-tree map,” wherein gene tree topologies are mapped to possible species tree topologies without assuming an underlying process. This was the approach taken by Ané et al. (2007), who used a Bayesian framework to estimate a species tree by maximizing gene tree concordance. Implemented in the software BUCKy (Larget et al. 2010), this method relies on the concept of concordance factors, or the proportion of gene trees for which a given bipartition is true (Baum 2007). The resulting phylogenetic estimate is referred to as the *primary concordance tree* (PCT), and can be estimated even if ILS is not the only process affecting gene tree incongruence. Larget et al. (2010) also introduced the concept of a population tree, which uses the average concordance factors for all quartets on an internal branch of the PCT to calculate branch lengths in coalescent units. Because concordance factors can contain information about both ILS and admixture/gene flow, Solís-Lemus and Ané (2016) developed a method for estimating species networks (species trees with reticulate edges) from concordance factors estimated for quartets of species. Their method, called SNaQ (Species Networks applying Quartets), uses these quartet concordance factors (QCFs) to maximize a pseudolikelihood function that matches the expected QCF values under the coalescent model with hybridization (Meng and Kubatko 2009) and the observed QCFs.

Despite the availability of these concordance factor-based approaches, there remain several areas where we believe the estimation of QCFs can be improved. First, estimating QCFs for multiple individuals or haplotypes per species is not easily accomplished using BUCKy or the methods available in the PhyloNetworks package that implements the SNaQ method (Ané et al. 2007; Solís-Lemus et al. 2017). This is problematic not only because having multiple alleles sampled from a population can potentially help to increase phylogenetic resolution (Andermann et al. 2018), but also because many hybrid plant lineages are polyploids, which means that not all of their homoeologs can be modeled simultaneously. Second, BUCKy requires the estimation of posterior distributions of gene trees, which can be computationally demanding for large numbers of loci and/or large numbers of sampled alleles. Using gene tree posteriors is a common way to deal with gene tree uncertainty (utilized by several methods in PhyloNet; Wen et al. 2018), but other methods, such as those using site pattern frequencies, would allow for faster computation of the gene tree quartet topologies that are used to estimate QCFs and could include measures of uncertainty from statistical resampling.

To address the issues listed above, we first developed a method for estimating concordance factors directly from sequence data for quartets of species. Our method accommodates multiple haplotypes sampled per species, and can conduct bootstrapping to account for gene tree uncertainty. To validate the method, we simulated multilocus sequence data on both tree and network topologies to assess how accurately it could estimate QCFs. We then collected nuclear amplicon data for two subsections in the plant genus *Penstemon* (Plantaginaceae; subsect. *Humiles* and *Proceri*). Subsections *Humiles* and *Proceri* are known to hybridize, and have the additional complication of containing multiple polyploid species. Using phased haplotypes from the nuclear amplicon sequences, we estimated species trees and networks using four different approaches to evaluate if the current circumscription of the subsections are in agreement with our phylogenetic estimates. Overall, we found strong evidence for hybridization in subsect. *Humiles* and *Proceri*, but phylogenetic support was generally lacking for many of the species relationships, owing to large amounts of genealogical discordance. Given the pace with which *Penstemon* has recently radiated, these types of patterns are not unexpected (Wolfe et al. 2006). Nevertheless, our phylogenetic estimates showed some stable relationships among the different methods used, and suggest that the current taxonomy of the two subsections needs revising.

## 2 Approach

We begin with a brief description of our method for QCF estimation, which uses site patterns to estimate the gene tree quartet relationships that are used to calculate species-level concordance factors. The basis for this method stems from ideas regarding the use of phylogenetic invariants for the inference of phylogenetic trees (Allman et al. 2008; Chifman and Kubatko 2015). For a given quartet of species, the QCF values represent the proportion of gene trees that agree with each of the three possible unrooted topologies relating the four species: ((1,2)(3,4)), ((1,3)(2,4)), and ((1,4)(2,3)). Because they represent a single bipartition among four species, these topologies are referred to as “splits”, and are denoted 12|34, 13|24, and 14|23 (Chifman and Kubatko 2015). Estimating the species-level concordance factors then amounts to estimating quartet topologies for each gene, followed by tabulating which species-level split is supported by each gene. When there are multiple haplotypes at a locus, we consider all of their possible sampling combinations and calculate the gene tree quartet topologies that support each species-level relationship. Using this approach, we are able to quickly estimate QCF values for samples with different ploidy levels. Below we detail our method for scoring gene tree quartet topologies, and combining them across sampled haplotypes to get species-level QCFs.

### Calculating Quartet Concordance Factors

Consider four species (1, 2, 3, and 4) with DNA sequence data collected at *G* independent loci, and with haplotypes phased and aligned for each locus. For example, a diploid individual from species 1 might have two haplotypes at locus one, which we denote 1_(1,1)_ and 1_(1,2)_. For any species, *S*, we denote its haplotypes at each gene by indexing across the gene number (*g* = 1, …, *G*) and the haplotype number (*h* = 1, …, *s*_*g*_; where *s*_*g*_ is the number of haplotypes present in species *S* at gene *g*). Then, for each gene, we score the three possible splits of all haplotype combinations using the frequency of matching site patterns. To get these scores, we first calculate the number of times a pair of species, *A* and *B*, have the same nucleotide at each site in the alignment:

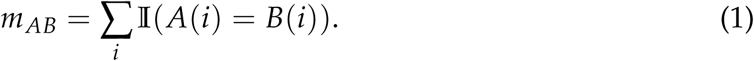

Here 𝕀() is the indicator function and is equal to 1 if the two bases are the same, and 0 otherwise.

We then use the *m*_*ab*_’s to calculate scores for each of the three unrooted quartet topologies:

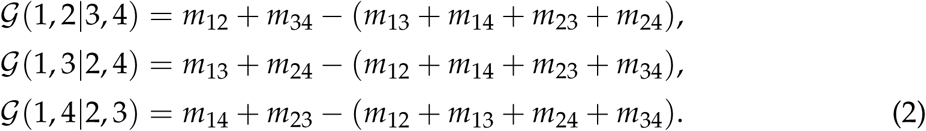

For these scores, patterns of nucleotide substitution that support a given split are given positive weight, while those that support alternative topologies are given negative weight. At the species level, we tabulate the number of gene trees supporting these same splits, and add 1 to the species topology that corresponds to the gene split with the highest score. If there is a tie for the highest score, we add 0.5 to the two species-level splits for the highest scoring gene trees. If all three gene tree splits have the same score, then 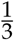 is added to each species-level split.

The calculation of concordance factors for each species-level quartet is then done by summing over all genes and tabulating how often each possible split is supported by each gene. This sum is also taken over all possible combinations of haplotypes, giving the following equation for calculating QCFs:

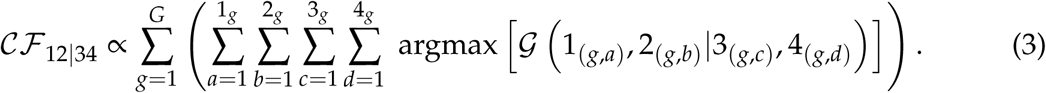

Here, argmax() is an indicator that is 1 if 𝒢(1, 2|3, 4) is the maximum argument and 0 otherwise. Ties are handled as described above. The calculation of 𝒞ℱ _13|24_ and 𝒞ℱ _14|24_ is the same as above but with the species and sums switched into the correct order for the split under consideration. All three species-level concordance factors are then normalized by their sum.

### Bootstrapping and Gene Tree Uncertainty

To deal with uncertainty in gene tree quartet estimation, we can also conduct bootstrap resampling of sites within genes when calculating the gene tree split scores. If we conduct *B* rounds of resampling, the gene tree contributions to the species-level splits can then be calculated across bootstrap replicates, with each gene tree split getting a weight proportional to the number of times it was the best scoring topology across all replicates:

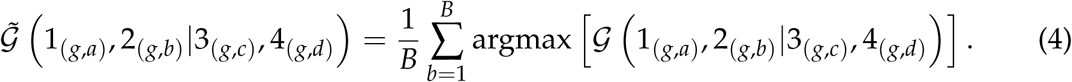

The species-level QCF value is then taken as the sum over these bootstrap weighted gene tree splits:

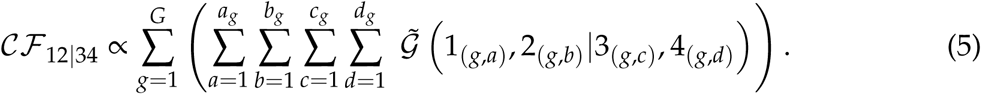

As before, 𝒞ℱ _13|24_ and 𝒞ℱ _14|23_ are calculated in a similar way, such that their indices are in the correct order. These 𝒞ℱ values are then normalized by their sum.

### Validating QCF Estimation

We validated our approach for QCF estimation using simulations on both tree and network topologies (Figure 1). The details of these simulations can be found in the online Supplemental Materials available on fig**share** (DOI: 10.6084/m9.figshare.11816682). In general, our method for calculating QCF values produced accurate estimates when compared to the true simulated data, with RMSD values ranging from 0.019–0.042 (0.023– 0.059 for bootstrapped) and 0.019–0.036 (0.021–0.043 for bootstrapped) for the tree and network topologies, respectively (Tables S4 and S5). Figures S1 and S2 show plots of fitted linear regression models for each quartet of species and the corresponding QCF estimate (x-axis) for each of the three unrooted topologies compared to the true simulated value (y-axis). Simulated QCFs were tabulated using PhyloNetworks (Solís-Lemus et al. 2017).

**Figure 1:**
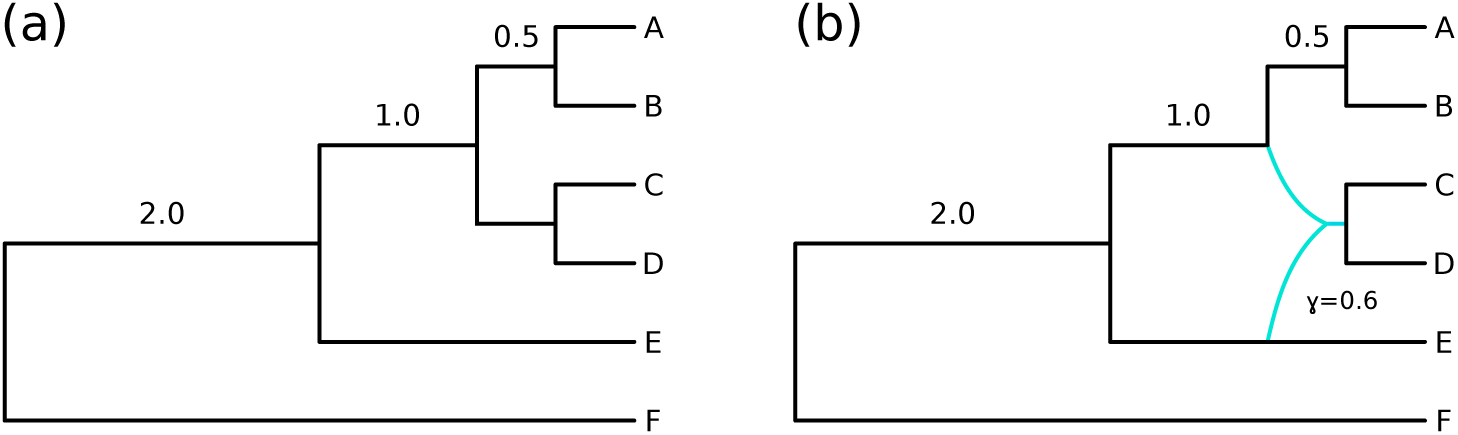
Simulation setup for (a) tree and (b) network topologies. Internal branches are annotated with their lengths in coalescent units (CUs). The total tree height is 4.0 CUs.

### Implementation

We have implemented our new method in the open-source software package `qcf`. `qcf` is wrtten in C++ and is available under the GNU GPL v3 on GitHub (https://github.com/pblischak/QCF). Documentation and tutorials for using the software can be found on ReadTheDocs (https://qcf.readthedocs.io).

## 3 Materials and Methods

### Study System

*Penstemon* Mitch. (Plantaginaceae) is the largest group of flowering plants endemic to North America, with ca. 300 species distributed from Alaska to Guatemala, and from the Pacific to Atlantic coasts (Wolfe et al. 2006). The center of diversity for *Penstemon* is the Intermountain West of the United States, with the biogeographic origin of the genus hypothesized to be in the Columbia Plateau (Straw 1966; Wolfe et al. 2002, 2006). *Penstemon* has undergone a recent and rapid radiation, which is thought to be driven by Pleistocene glaciation cycles, as well as adaptation to different ecological habitats and pollinators (Castellanos et al. 2006; Wolfe et al. 2006; Wilson et al. 2007). The most comprehensive molecular phylogeny of *Penstemon* was published by Wolfe et al. (2006), with 193 species sampled for their analyses of the nuclear ribosomal ITS region and two chloroplast genes (*trnCD*+*trnTL*). While support for many species-level relationships was lacking, Wolfe et al. (2006) were able to make several inferences regarding higher level relationships within *Penstemon*. A more recent study conducted high-throughput sequencing for 70 species of *Penstemon* (Wessinger et al. 2016), and recovered high support across the entire tree. However, the limited taxon sampling of their phylogeny did not contain many members of the subg. *Penstemon*, limiting the interpretation of their results in the context of the whole genus.

Two groups within *Penstemon* subg. *Penstemon* that are of particular interest are the subsections *Humiles* and *Proceri*. Species in these subsections are primarily distributed in the Pacific Northwest of the United States, and occur at subalpine to alpine elevations in a variety of habitats, including some that are atypical for species of *Penstemon* in the western US (e.g., mesic meadows; Keck 1945; Nold 1999). These subsections are also morphologically distinct from other members of the genus, with their inflorescences organized into verticillasters. Many of the species also have glandular hairs on the inflorescence, a character present in all species of subsect. *Humiles*, but only in some members of subsect. *Proceri* (Keck 1945). The traditional taxonomic division between these groups is based on a single leaf character: members of subsect. *Humiles* have serrate leaf margins and members of subsect. *Proceri* have entire margins (Keck 1945). However, there have been observations of hybridization between the two subsections, such that members of subsect. *Proceri* can sometimes have toothed leaf margins (Strickler 1997). Cases of hybridization at the diploid level are well documented in *Penstemon* (Straw 1955; Crosswhite 1965; Wolfe et al. 1998b,a; Datwyler and Wolfe 2004), and numerous instances of polyploidy, mostly within sect. *Penstemon* and subg. *Saccanthera*, have been studied as well (Keck 1945; Broderick et al. 2011). Most of these polyploids are thought to be allopolyploids (formed through hybridization) (Wolfe et al. 2006), but the majority of these hypotheses remain untested (but see Lawrence and Datwyler 2016).

Given the lability of the leaf character dividing these two subsections, as well as their similarities in geographic ranges and morphology, we sought to evaluate the monophyly of subsect. *Humiles* and *Proceri* using nuclear amplicon data. We also aimed to investigate the extent to which hybridization has occurred in these groups, as well as gaining an understanding of the origin of any polyploid taxa (auto-vs. allopolyploid). In the case of subsect. *Humiles* and *Proceri*, the *P. attenuatus* species complex presents a compelling test for understanding polyploidy in these groups. According to Keck (1945), the four varieties of *P. attenuatus* are all hypothesized to be allopolyploids, forming through hybridization between *P. albertinus* in subsect. *Humiles* and three different species in subsect. *Proceri* (see Figure 2). Earlier molecular phylogenetic analyses recovered the subsections as polyphyletic (Wolfe et al. 2006). However, these patterns are based on two uncombined gene trees, which would not allow for processes that cause gene tree incongruence to be modeled. Here we use species tree and network approaches to account for genealogical discordance caused by both ILS and hybridization to estimate a phylogeny for subsect. *Humiles* and *Proceri*.

**Figure 2:**
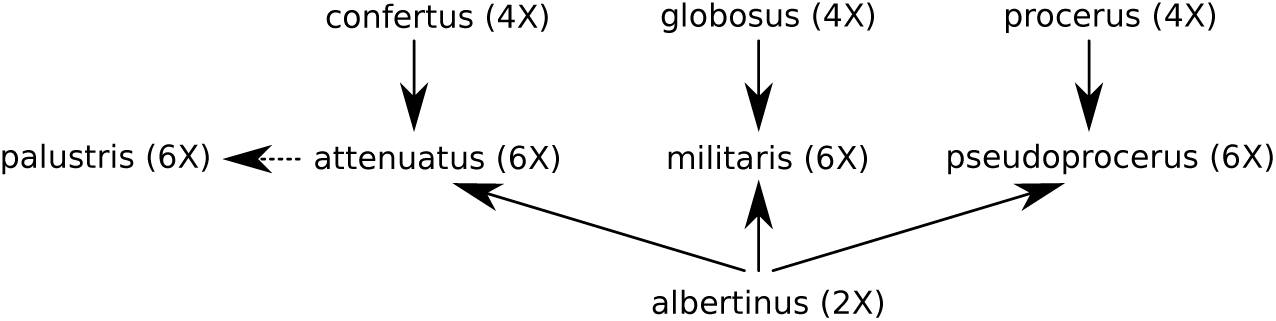
Hypotheses of allopolyploid formation in *Penstemon attenuatus* according to Keck (1945). Varieties of *P. attenuatus* are placed in the center, with their putative diploid parent below (*P. albertinus* for all), and their putative tetraploid parents above. *P. attenuatus* var. *palustris* is marked with a dashed arrow due to the uncertainty of its placement.

### Sample Collection, DNA Extraction, and Amplicon Sequencing

DNA was extracted from field-collected leaf tissue that was dried on silica gel. We used a modified CTAB protocol for DNA isolation (Wolfe 2005), quantified all samples using a Qubit fluorometer (Invitrogen, Carlsbad, CA, USA), and normalized all samples to 20 ng/*µ*L. Normalized DNA samples for 38 accessions representing 17/22 and 20/27 currently circumscribed taxa from subsect. *Humiles* and *Proceri*, respectively, plus an out-group taxon from *Penstemon* subgenus *Dasanthera* (*P. davidsonii* var. *davidsonii*), were sent to the IBEST Genomics Resources Core at the University of Idaho (Moscow, ID, USA) for sample preparation and sequencing (listed in Table S1). Amplification of targeted amplicons and the addition of sample barcodes and Illumina adapters were done using microfluidic PCR on the Fluidigm 48 x 48 Access Array (Fluidigm Corporation, South San Francisco, CA, USA), followed by 300 bp paired-end sequencing on an Illumina MiSeq (Illumina, San Diego, CA, USA) (Uribe-Convers et al. 2016). Primers for the 48 loci used in this study were designed and tested as described in Blischak et al. (2014), and are given in Tables S2 and S3.

Raw, paired-end sequencing reads were returned from IBEST and processed using Fluidigm2PURC v0.1.2 (https://github.com/pblischak/fluidigm2purc; Blischak et al. 2018b). Fluidigm2PURC trims reads using Sickle (Joshi and Fash 2011), joins paired reads using FLASH2 (Magoč and Salzberg 2011), and prepares the input file for clustering and chimera detection using the program PURC (Rothfels et al. 2017). After clustering with PURC, haplotypes are inferred based on cluster sizes and user-specified ploidy levels. Three rounds of chimera detection and clustering were performed using default settings in the script `purc recluster2.py`, a modified version of the original script distributed with PURC (https://bitbucket.org/crothfels/purc). To get haplotypes, clusters were first cleaned of excessive gaps using Phyutility (threshold = 33%; Smith and Dunn 2008) and then realigned using MAFFT (Katoh 2013). Haplotypes were then inferred for all sampled taxa assuming known ploidy levels reported in Keck (1945), Strickler (1997), and Broderick et al. (2011). Information regarding haplotype dosage (i.e., number of haplotype copies) was ignored, resulting in only unique haplotypes being returned for each taxon at each gene.

### Species Tree Inference

Haplotype-level gene trees for each locus were inferred with RAxML v8.2.11 using the GTRGAMMA model of nucleotide substitution and 500 rapid bootstrap replicates (Stamatakis et al. 2008; Stamatakis 2014). We then inferred a taxon-level species tree with ASTRAL v5.5.9 (ASTRAL-III) using a mapping file to link haplotypes with their respective taxa (Mirarab and Warnow 2015). To increase the thoroughness of the ASTRAL-III search algorithm, we added the following command line options: `--polylimit 20` (maximum size of polytomy), `--samplingrounds 100` (number of rounds of subsampling haplotypes from taxa), and `--extraLevel 2` (increase the number of bipartitions added to the search space).

A species tree was also inferred using methods from the TICR pipeline (https://github.com/nstenz/TICR; Stenz et al. 2015). The TICR pipeline estimates QCFs using BUCKy (Larget et al. 2010), and then uses the concordance values of these quartets to infer a species tree using the QuartetMaxCut algorithm (Snir 2012). Here, we instead used our new method to estimate QCFs, and inferred a species tree using the script `get-pop-tree.pl`. Average concordance factors and branch lengths (in coalescent units) were then estimated for this tree using the `getTreeBranchLengths.R` script.

As a final estimate of phylogeny, we used only the majority haplotype (haplotype inferred from the largest cluster) returned by Fluidigm2PURC to estimate a species tree with RAxML using a supermatrix as input. It has been shown previously that concatenating multilocus data can result in incorrect inferences of phylogeny when gene tree discordance is present (Kubatko and Degnan 2007). However, congruence among different methods can also be a good indicator of stable species relationships. Inference with RAxML was conducted using a partition file to estimate separate model parameters for each gene, and 1000 rapid bootstrap replicates were used to assess statistical support.

### Candidate Hybridization Events from Rooted Triples

To generate a list of candidate hybridization events, we used the program HyDe v0.4.2 to test for hybridization on all possible triples of taxa from subsect. *Humiles* and *Proceri* (Blischak et al. 2018a). HyDe tests for hybridization using site pattern frequencies (Kubatko and Chifman 2015), and estimates the amount of admixture occurring between two parental taxa to form a third hybrid taxon. Using *P. davidsonii* var. *davidsonii* as an out-group and a mapping file to assign haplotypes to taxa, we tested all triples in all directions using the `run_hyde_mp.py` script. Statistical significance was assessed at the *α* = 0.05 level with a Bonferroni correction for the number of hypothesis tests conducted.

### Species Network Inference

Our species tree analyses with ASTRAL-III, QuartetMaxCut, and RAxML (supermatrix) recovered two clades with corresponding taxon membership (see Results), which we refer to as clades A and B (Figures 3–5). To reduce the computational burden of estimating a large network (*>* 30 taxa), we chose to analyze these clades independently. Haplotypes from taxa belonging to each clade were extracted from the original sequence alignments and written to new files. *Penstemon davidsonii* var. *davidsonii* was included in the data set for both clades as an outgroup. We then estimated haplotype-level gene trees as before, and inferred a taxon-level species in ASTRAL-III using a mapping file and default search settings. We also estimated QCFs for each clade using the `qcf` software with 500 bootstrap replicates. The resulting species trees and QCF estimates for clades A and B were then used as input for network estimation using the SNaQ method implemented in the software package PhyloNetworks v0.7.0 (Solís-Lemus and Ané 2016; Solís-Lemus et al. 2017). We varied the maximum number of hybridization events from h=1 to h=5 and used the resulting log-pseudolikelihood values to determine the most likely number of hybridization events. The log-pseudolikelihood for the case of no hybridization (h=0) was calculated by maximizing the fit of the observed QCF values on the fixed tree topology estimated by ASTRAL-III. All network analyses were conducted on the Oakley cluster at the Ohio Supercomputer Center (https://www.osc.edu).

**Figure 3:**
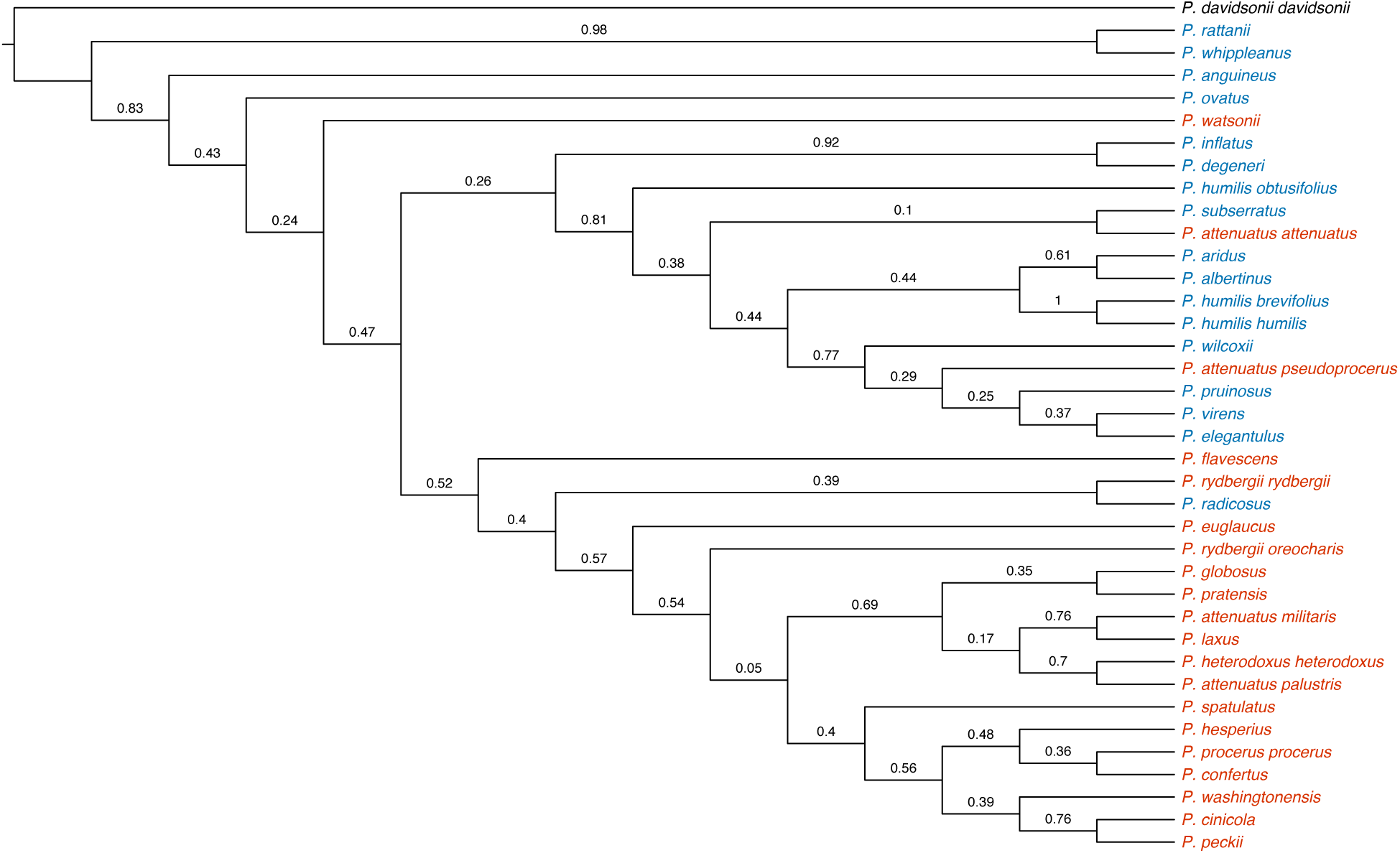
Phylogeny of *Penstemon* subsections *Humiles* and *Proceri* inferred by ASTRAL-III. Labels on branches are local posterior probabilities (Sayyari and Mirarab 2016). Taxa are colored based on their current taxonomic classification: red = subsect. *Proceri*, blue = subsect. *Humiles*.

**Figure 4:**
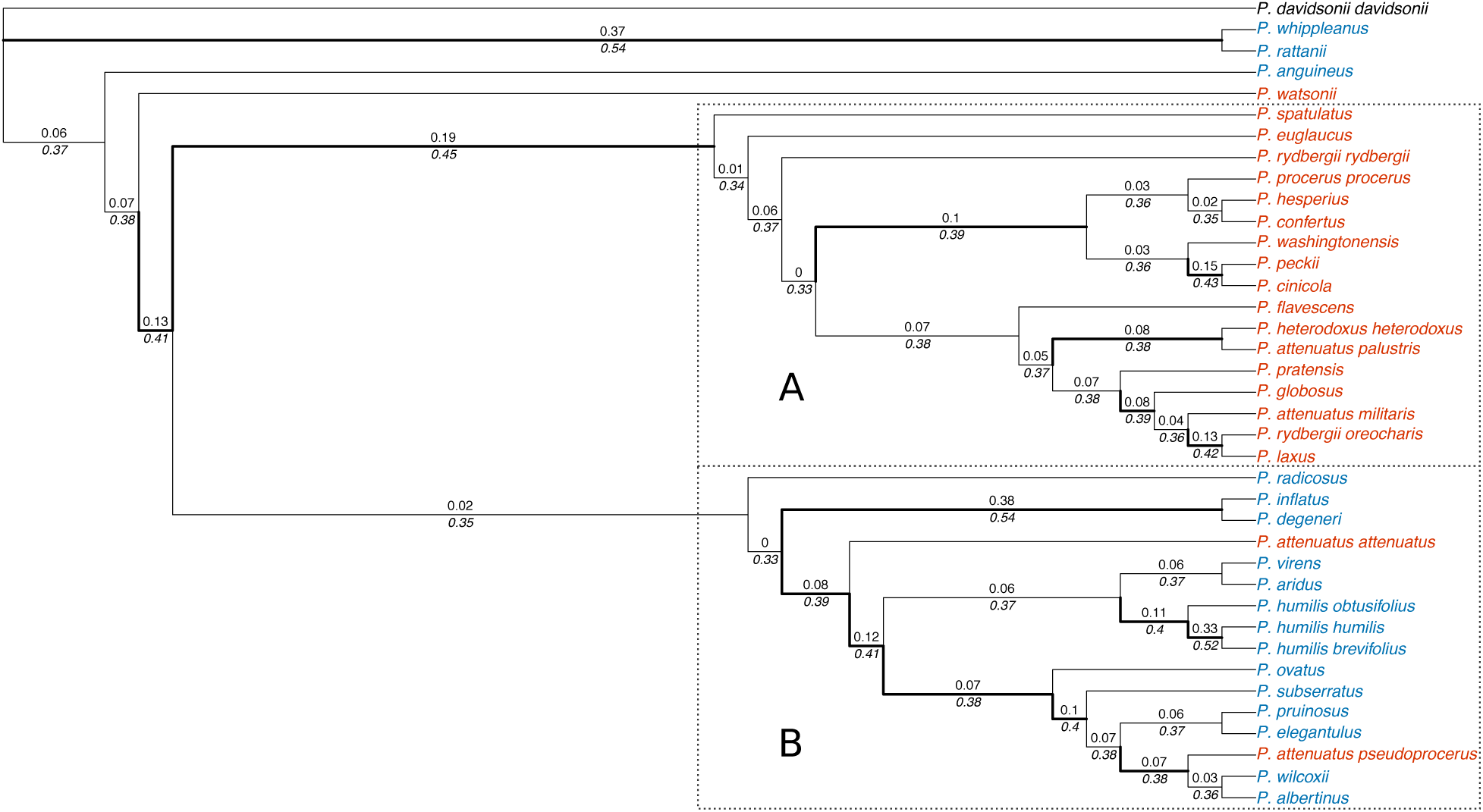
Phylogeny of *Penstemon* subsections *Humiles* and *Proceri* inferred using qcf and Quartet-MaxCut. Each branch is labeled above by its length in coalescent units and below by the average QCF value for all quartets induced by that branch. All branches with average QCF values greater than 0.38 are plotted with thicker lines (Stenz et al. 2015). Clades outlined by boxes labeled ‘A’ and ‘B’ correspond to the groups that were used for species network inference. Taxa are colored the same as in Figure 3.

**Figure 5:**
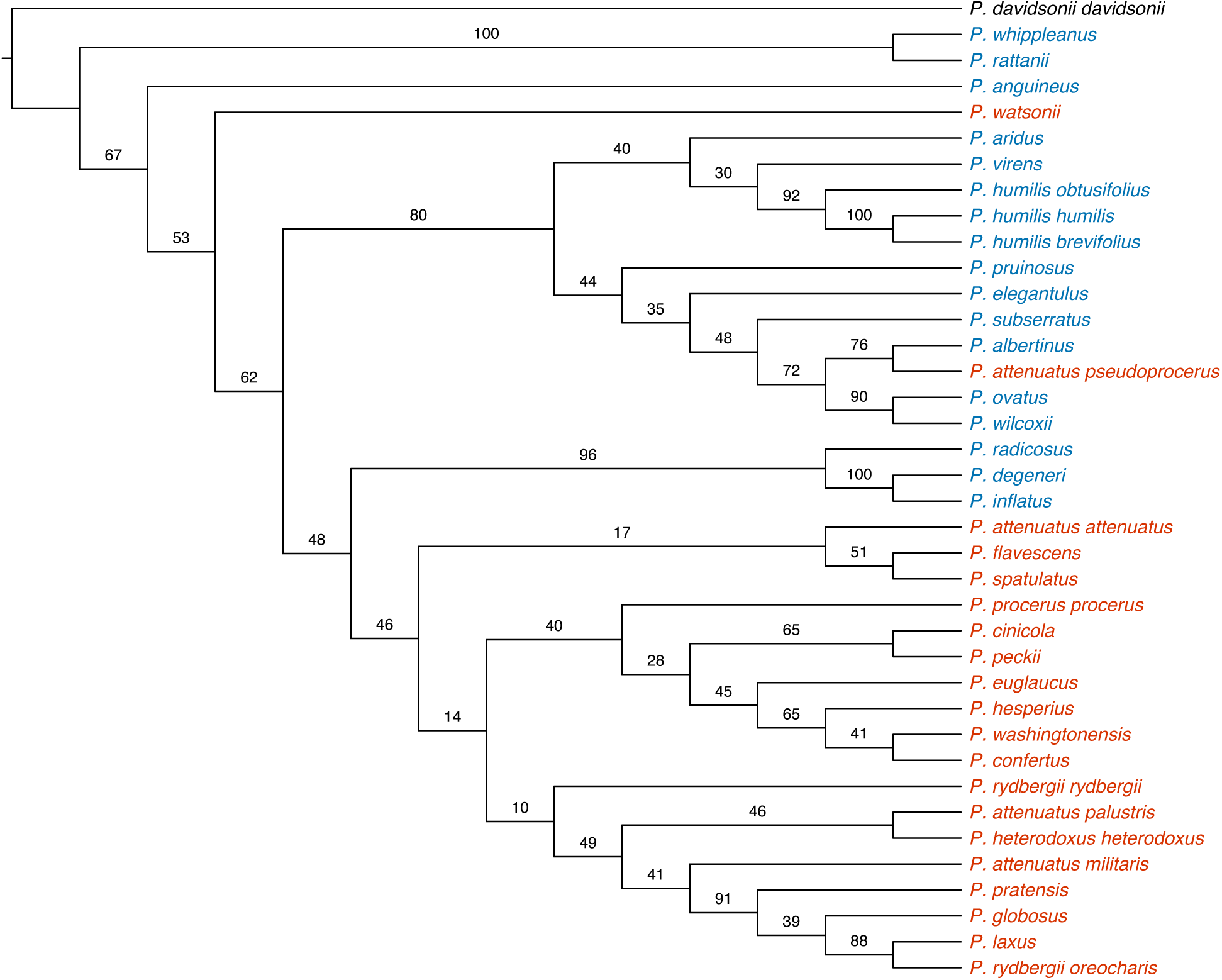
Phylogeny of *Penstemon* subsections *Humiles* and *Proceri* inferred with RAxML using a supermatrix of 43 loci. Labels on branches are support values from 1000 bootstrap replicates. Taxa are colored the same as in Figure 3.

## 4 Results

### Nuclear Amplicon Data

Of the 48 loci that were amplified using the Fluidigm Access Array, 43 (all nuclear) recovered sufficient data for processing and downstream phylogenetic analyses. Data processing with Fluidigm2PURC on these 43 loci produced phased haplotypes for all 38 taxa, with many of the polyploid taxa containing three or more unique haplotypes. The supermatrix of majority haplotypes used for analysis with RAxML had a total alignment length of 18,207 bp.

### Species Tree Inference

Species trees were inferred with three different methods, all of which produced mostly similar phylogenetic estimates for subsect. *Humiles* and *Proceri* (Figures 3–5). For each method, four species were consistently recovered as a grade outside of the rest of the ingroup: *P. anguineus, P. rattanii, P. watsonii*, and *P. whippleanus. Penstemon ovatus* was also recovered outside of the two subsections in the ASTRAL-III analysis. For ASTRAL-III and QuartetMaxCut, two species from subsect. *Proceri, P. attenuatus* var. *attenuatus* and *P. attenuatus* var. *pseudoprocerus*, were recovered in a clade consisting primarily of species from subsect. *Humiles*. These two methods also recovered a clade consisting almost entirely of species from subsect. *Proceri*, with the exception of *P. radicosus* being present in the ASTRAL-III tree. The supermatrix analysis inferred a tree with a number of relationships that differed from the other two approaches. Three of the four variaties of *P. attenuatus* were inferred to belong to the same clade, but *P. attenuatus* var. *pseudoprocerus* was still recovered in a clade of *Humiles* taxa. This analysis also shifted a clade of three species, *P. radicosus, P. degeneri*, and *P. inflatus*, to be sister to the clade consisting of *Proceri* species with high support (bootstrap support = 96%). Another notable difference among the methods was that ASTRAL-III did not recover the three varieties of *P. humilis* as monophyletic, but the QuartetMaxCut and supermatrix approaches did.

Estimated branch lengths in coalescent units from ASTRAL-III and QuartetMaxCut were extremely short for all internal branches, indicating rampant genealogical discordance (Figures 4, S3, and S4). Branch lengths from the RAxML supermatrix analysis were also very short for the branches along the backbone of the tree, demonstrating that few substitutions were present to inform relationships for these deeper bipartitions. Support values were generally low across the different trees, with only a few relationships showing high levels of support. This is likely a result of the short branches observed in the different trees, and support the hypothesis that speciation has occurred rapidly, with little time for informative substitutions to occur. Another possible reason for these low support values is the occurrence of hybridization. As we show below, there is strong evidence for hybridization in these groups, making phylogenetic inference difficult.

### Tests for Hybridization and Species Network Inference

Analyzing all possible triples of ingroup taxa with HyDe resulted in a total of 23,310 hypothesis tests, of which 282 showed significant evidence for hybridization. The average value for the hybridization parameter (*γ*) was 0.513 (standard deviation = 0.114), with a minimum and maximum value of 0.205 and 0.843, respectively. Out of 37 total ingroup taxa, 24 had a significant signal for hybridization.

Species network inference with SNaQ was then conducted on the two primary clades that were recovered in the QuartetMaxCut analyses (clades A and B; Figure 4). The reason for using these clades was that this analysis recovered a pattern of relationships for subsect. *Humiles* and *Proceri* that was most consistent with the current taxonomy. Although hybridization was detected between species in these clades using HyDe, network inference with SNaQ cannot currently handle 38 taxa. However, since the members of these clades were recovered fairly consistently between the different methods that we used for species tree inference, we decided to analyze them independently to make network inference computationally feasible.

Using a range of values on the number of possible reticulation events (h=0 to h=5), we were able to infer species networks in all cases for both clades A and B within the amount of compute time allotted (10 cores, 80–100 hours). For both clades, adding reticulation events greatly reduced the log-pseudolikelihood, providing strong evidence that hybridization is occurring within these clades. Networks with four and three reticulations had the highest pseudolikelihoods for clades A and B, respectively, but the network topology with four reticulations for network A produced non-sensical relationships (hybridization with the outgroup), so we preferred the network with h=3 (Figures S5 and S6). In addition to having the highest (or second highest for clade A) log-pseduolikelihood, the networks estimated with three reticulations were among the only estimates that had sensible branch lengths. For most other networks in both clades A and B, one of the reticulate edges was always inferred to have a branch length of *∼*9.5 coalescent units. Given the amount of gene tree discordance present in the data set, and the short branch lengths estimated by ASTRAL-III and QuartetMaxCut, these estimates are most likely incorrect.

The best networks for clades A and B showed different patterns for the timing of hybridization (Figure 6). For clade A, all of the reticulation events occurred closer to the present, and only involved pairs of species hybridizing. The hybridization events inferred include: (1) *P. spatulatus* × *P. attenuatus* var. *palustris* → *P. flavescens*, (2) *P. rydbergii* var. *oreocharis* × *P. globosus* → *P. laxus*, and (3) *P. confertus* × *P. cinicola* → *P. washingtonensis*. Clade B, on the other hand, was estimated to have a deep reticulation event involving two ancestral populations, with the resulting hybrid lineage then diversifying into 12 different taxa. The other hybridization event in this clade was *P. ovatus* × *P. humilis* var. *humilis* → *P. humilis* var. *brevifolius*.

**Figure 6:**
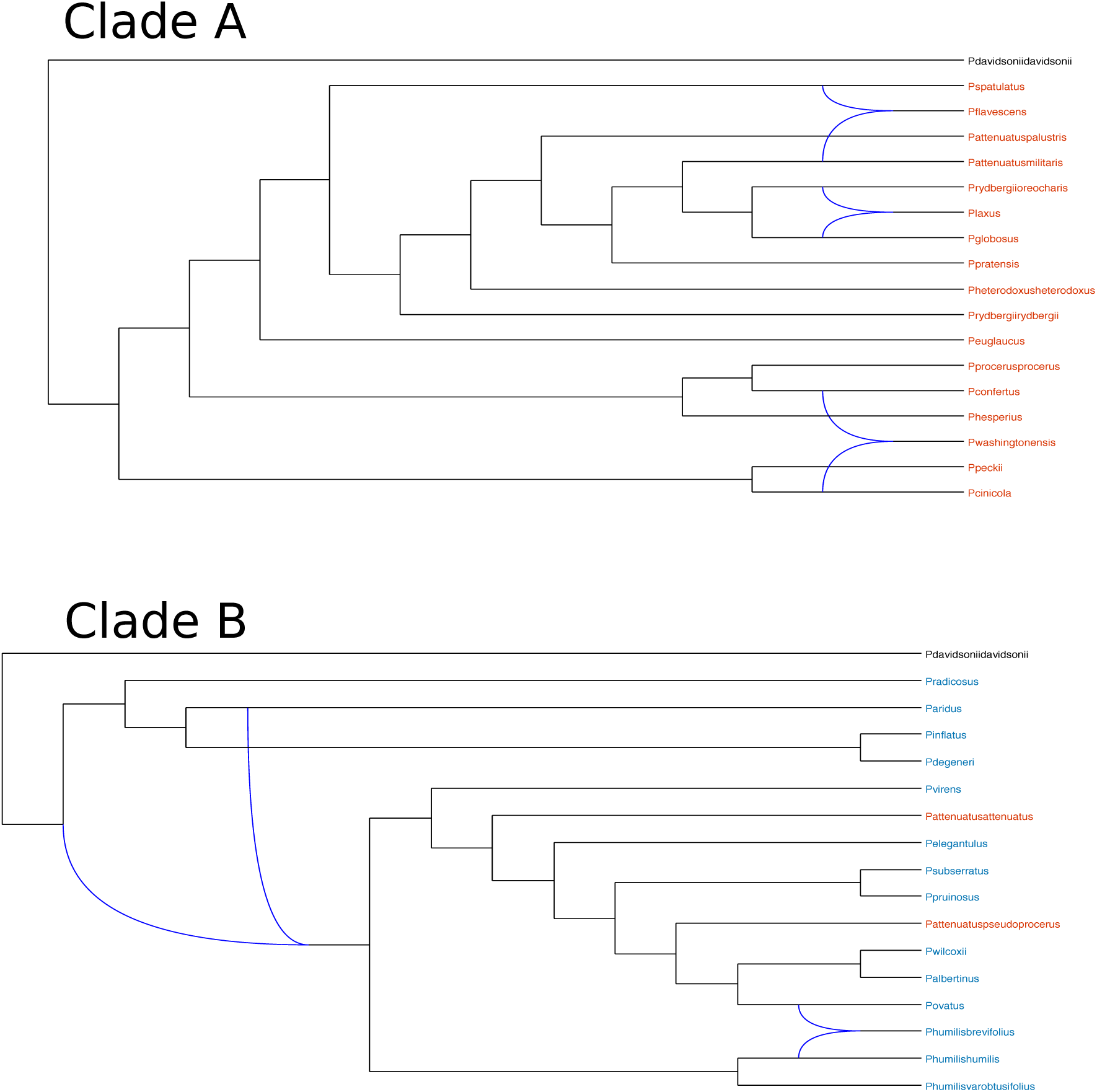
Best maximum pseudolikelihood (ML) networks for clades A and B estimated by SNaQ. The maximum number of hybridization events for these ML networks is h=3. Taxa are colored the same as in Figure 3.

## 5 Discussion

In this paper, we investigated the phylogenetic relationships and taxonomic affinities among the taxa within *Penstemon* subsections *Humiles* and *Proceri* using nuclear amplicon sequencing. We found strong evidence for hybridization in these groups, but the rapid diversification in these two subsections made the exact inference, localization, and interpretation of reticulation events extremely difficult. Despite these shortcomings, there are some clear trends regarding the taxonomic implications of our phylogenetic estimates, as well as what they may mean for character evolution and biogeography in the group. Our work also highlights the difficulties of estimating phylogeny for recently radiating groups with variable ploidy levels and high amounts of hybridization, an issue that is common for many groups of angiosperms.

### Taxonomy of Subsections *Humiles* and *Proceri*

Using several methods for phylogenetic inference, we found evidence supporting the non-monophyly of subsect. *Humiles* and *Proceri* (Figures 3–5). This pattern was recovered for all methods, despite variable levels of statistical support for the different analyses, suggesting that this pattern is robust to the various assumptions of each method. Four species in particular were recovered completely outside of the two subsections: *P. whippleanus, P. rattanii*, and *P. anguineus* (all subsect. *Humiles*), as well as *P. watsonii* (subsect. *Proceri*). Only one of the methods, QuartetMaxCut, recovered a monophyletic grouping of species belonging to subsect. *Proceri*. However, this analysis also placed two taxa currently classified in subsect. *Proceri* (*P. attenuatus* var. *attenuatus* and *P. attenuatus* var. *pseudoprocerus*) into a clade of species from subsect. *Humiles*, casting doublt on Keck’s hypotheses of allopolyploid formation for these taxa (Keck 1945). Interestingly, none of the varieties of *P. attenuatus* showed the predicted affinities for their putative parental taxa (Figure 2). However, given their hypothesized hybrid nature, it is possible that they are simply difficult to place using models that do not include reticulation.

The phylogenetic placement of hybrid taxa has been shown to be problematic in phylogenetic analyses, with hybrids typically branching at the base of a clade containing one of their parental taxa (e.g., McDade 1990, 1992). Our tests for hybridization and species network analyses confirmed the presence of hybrids within and between subsect. *Humiles* and *Proceri*, but did not support the putative parentage of the varieties of *P. attenuatus*, as well as a number of other hypothesized hybrids from Keck (1945). Nevertheless, the occurrence of a deep hybridization event in the clade consisting of taxa primarily from subsect. *Humiles* (clade B) is especially interesting. The impact of hybridization on genetic variation and its connection to the subsequent speciation and diversification of hybrid lineages has long been understood in plants (Anderson 1949; Stebbins 1950; Anderson and Stebbins 1954; Grant 1971; Mallet 2007), and several recent studies have observed deep hybridization events at the diploid (e.g., Folk et al. 2016; García et al. 2017; Folk et al. 2018) and polyploid (e.g., Morales-Briones et al. 2018) levels. The hybrid group within clade B is a mix of diploids and polyploids, suggesting that the processes of both hybridization and whole genome duplication have been at play in its diversification. Future work with more genomic data will be important for this clade to gain better resolution for any additional hybridization events.

A particular strength of our method of QCF estimation is the ability to analyze taxa with different ploidy levels, allowing us to analyze all diploid and polyploid taxa in subsect. *Humiles* and *Proceri* simultaneously. From our network analyses with SNaQ, the only polyploid that was inferred to be a hybrid was *P. flavescens* (hexaploid; clade A). However, our tests for hybridization with HyDe found far more evidence for hybridization, likely because it only tests three species at a time, rather than trying to infer an entire network. Nevertheless, out of the 24 taxa inferred to be hybrids using HyDe, only four were polyploids. This casts doubt on the hypothesis that most of the polyploids in subsect. *Proceri* are of hybrid origin. However, a lack of phylogenetically informative variation could also be preventing us from detecting the full extent of hybridization potentially occurring in these polyploids.

### Character Evolution and Biogeography

Given the reticulate history of subsect. *Humiles* and *Proceri*, there are several patterns of morphological character evolution that can be interpreted in the context of their past genetic exchanges. Of particular interest is the presence of glandular hairs on the inflorescence, a trait with potentially adaptive importance (Levin 1973) that is present in all species of subsect. *Humiles*, but is absent in the majority of species in subsect. *Proceri*. For the species in subsect. *Proceri* where it does occur, it is hard to determine if it is simply a labile trait that has arisen several times, or if there is a single clade of species that all have the trait. A perhaps more interesting scenario could be that this trait was gained through hybridization or introgression, however testing this hypothesis is currently not feasible due to a lack of methods for discrete character reconstruction on phylogenetic networks (but see Jhwueng and O’Meara 2015; Bastide et al. 2018, for examples of continuous character evolution). A possible workaround would be to construct all possible resolutions of the underlying trees displayed by the networks and to reconstruct the character history on each tree. Nevertheless, future model development on discrete character evolution on networks will help to address this type of question.

The biogeographic context of these hybridization events is also of interest, with reconstructions of species’ geographic ranges potentially helping to shed light on the plausibility of hypotheses about the occurrence of reticulation. The current geographic distribution of the taxa in subsect. *Humiles* and *Proceri* is concentrated in the Pacific Northwest of the United States, an area with several well-established biogeographic and phylogeographic hypotheses regarding the occurrence of species in the Cascade Range, the Northern Rocky Mountains, and the Sierra Nevada in northern California (Brunsfeld et al. 2001; Carstens et al. 2005; Brunsfeld et al. 2007). Two recent studies that have investigated the biogeography of hybridization events include Burbink and Gehara (2018) and Folk et al. (2018), who take different approaches to reconstructing ancestral contact zones where hybridization could potentially have occurred. Burbink and Gehara (2018) found a single deep reticulation event in the phylogeny of New World kingsnakes and used the resulting parental trees (trees where a hybrid clade is sister to either parent) to infer ancestral areas for the hybrid clade. Folk et al. (2018) used climatic niche reconstructions to find likely regions where ancestral lineages of *Huechera* and *Mitella* may have occurred in sympatry and hybridized. These approaches could be used in concert to illuminate the dynamics of vicariance and dispersal for lineages of subsect. *Humiles* and *Proceri*, as well as helping to locate geographic regions where hybridization, as well as whole genome duplication, could have occurred in the past.

### Phylogenetics of Hybrids and Polyploids

As was seen from our phylogenetic analyses, the internal branch lengths of our species trees and networks were very short (most were less than 0.5 coalescent units), highlighting the prevalence of incomplete lineage sorting and genealogical discordance in our data. Previous research has shown that *Penstemon* is a young genus (crown age 2.5– 4.0 mya) that has radiated extremely rapidly, with hybridization and polyploidy occurring frequently (Wolfe et al. 2006; Wessinger et al. 2016). These types of processes are likely not uncommon for other groups of angiosperms, and having methods to deal with them will be especially important for making future inferences about the evolutionary history of these groups. To resolve hybridization events, especially when they involve polyploids, there are several methods that have already been developed. Some of these are not coalescent-based, but instead try to reconstruct a network from gene trees that have all of the haplotypes from a polyploid sampled (a so called “multi-labeled” tree; Lott et al. 2009; Marcussen et al. 2012). Other studies have relied on coalescent-based assignment of homoeologous haplotypes into putative, diploid subgenomes, but these approaches can be computationally limited due to the cost of exploring all permutations of haplotypes assignments (Bertrand et al. 2015; Oberpieler et al. 2017). The only approach to simultaneously infer a network topology and homoeolog assignment in a coalescent framework is the method of Jones et al. (2013). However, this method uses a hierarchical Bayesian framework that does not scale well to large numbers of loci or taxa.

If homoeolog assignment is the goal, then it may be beneficial to first identify parental taxa so that the number of comparisons for determining haplotype origin is reduced. Kamneva et al. (2017) used such an approach in strawberries to identify potential parents for several different polyploid species. They used a two-step approach to generate and test hypotheses, first constructing networks using consensus methods, followed by evaluating the likelihood of candidate networks using PhyloNet (Wen et al. 2018). Their analyses were limited to no more than 5 haplotypes per taxon, and also did not include an actual search over network space. Our method for QCF estimation was able to analyze all inferred haplotypes for all 38 taxa sampled in this study, and our network analyses with SNaQ were used to conduct an actual search over network topologies. The appeal of these types of approaches is that they do not require *a priori* knowledge about parental taxa when inferring a network. For non-model taxa where cases of hybridization and allopolyploidy are being investigated for the first time, the ability to model these processes with little input from the user regarding putative hybridization events should help to facilitate the discovery of reticulate evolutionary events in virtually any group of taxa where they may be occurring.

### 6 Conclusions

Hybridization and polyploidy are processes that obscure phylogenetic inference for many groups of taxa, and are a particular problem for lineages of angiosperms, where they are especially common. Using the concept of quartet concordance factors (the proportion of gene tree quartets supporting a species-level quartet), we developed a method for estimating these concordance factors that can accommodate taxa with variable ploidy levels. Using this approach, and several others, we then inferred species trees for *Penstemon* subsect. *Humiles* and *Proceri*, finding that the subsections were not reciprocally monophyletic. Tests for hybridization and species network inference also revealed that reticulation has been a common occurrence in these groups. In general, this study highlights the difficulties of inferring phylogeny in a rapid species radiation where hybridization and WGD are common. However, our approach for QCF estimation, in combination with the network inference method SNaQ, helped to disentangle the complex patterns of hybridization in these subsections, and should provide a useful tool for other researchers interested in reticulate evolution as well.

## Acknowledgements

The authors thank members of the Wolfe and Kubatko labs for comments that helped to improve previous versions of the manuscript. PDB would like to thank Mikel Stevens, Noel Holmgren, Karen and Steve Shelly, Carol Blackburn, Teresa Prendusi, Dale Reinhart, Maret Pajutee, and Steve Popovich for assistance in the field and for help with obtaining permits. This work was supported by the following grants: National Science Foundation DEB-1601096 to ADW and PDB (Doctoral Dissertation Improvement Grant), National Science Foundation DEB-1455399 to ADW and LSK, and Graduate Student Research Grants from the Society of Systematic Biologists and the American Society of Plant Taxonomists. Computational resources for this research were provided by the Ohio Supercomputer Center and the College of Arts and Sciences Unity Cluster at The Ohio State University.

## References

Allman, E. S., C. Ané, and J. A. Rhodes. 2008. Identifiability of a Markovian model of molecular evolution with Gamma-distributed rates. Advances in Applied Probability 40:229–249.

Andermann, T., A. M. Fernandes, U. Olsson, M. Töpel, B. E. Pfeil, B. Oxelman, A. Aleixo, B. C. Faircloth, and A. Antonelli. 2018. Allele phasing greatly improves the phylogenetic utility of ultraconserved elements. Systematic Biology https://doi.org/10.1093/sysbio/syy039.

Anderson, E. 1949. Introgressive hybridization. John Wiley, New York, NY, USA.

Anderson, E. and G. L. Stebbins. 1954. Hybridization as an evolutionary stimulus. Evolution 8:378–388.

Ané, C., B. Larget, D. A. Baum, and A. Rokas. 2007. Bayesian estimation of concordance among gene trees. Molecular Biology and Evolution 24:412–426.

Bastide, P., C. Solís-Lemus, R. Kriebel, K. W. Sparks, and C. Ané. 2018. Phylogenetic comparative methods on phylogenetic networks with reticulations. Systematic Biology https://doi.org/10.1093/sysbio/syy033.

Baum, D. A. 2007. Concordance trees, concordance factors, and the exploration of reticulate genealogy. Taxon 56:417–426.

Bertrand, Y. J. K., A.-C. Scheen, T. Marcussen, B. E. Pfeil, F. de Sousa, and B. Oxelman. 2015. Assignment of homoeologues to parental genomes in allopolyploids for species tree inference, with an example from *Fumaria* (Papaveraceae). Systematic Biology 64:448–471.

Blischak, P. D., J. Chifman, A. D. Wolfe, and L. S. Kubatko. 2018a. HyDe: a Python package for genome-scale hybridization detection. Systematic Biology https://doi.org/10.1093/sysbio/syy023.

Blischak, P. D., M. Latvis, D. F. Morales-Briones, J. C. Johnson, V. S. Di Stilio, A. D. Wolfe, and D. C. Tank. 2018b. Fluidigm2PURC: automated processing and haplotype inference for double-barcoded PCR amplicons. Applications in Plant Sciences 6:e1156.

Blischak, P. D., A. J. Wenzel, and A. D. Wolfe. 2014. Gene prediction and annotation in *Penstemon* (Plantaginaceae): a workflow for marker development from low-coverage genome sequencing. Applications in Plant Sciences 2:1400044.

Broderick, S. R., M. R. Stevens, B. Geary, S. L. Love, E. N. Jellen, R. B. Dockter, S. L. Daley, and D. T. Lindgren. 2011. A survey of *Penstemon*’s genome size. Genome 54:160–173.

Brunsfeld, S. J., T. R. Miller, and B. C. Carstens. 2007. Insights into the biogeography of the Pacific Northwest of North America: evidence from the phylogeography of *Salix melanopsis* (Salicaceae). Systematic Botany 32:129–139.

Brunsfeld, S. J., J. Sullivan, D. S. Soltis, and P. S. Soltis. 2001. Integrating ecological and evolutionary processes in a spatial context chap. Comparative phylogeography of northwestern North America: a synthesis, Pages 319–339. Oxford: Blackwell Science.

Burbink, F. T. and M. Gehara. 2018. The biogeography of deep time reticulation. Systematic Biology https://doi.org/10.1093/sysbio/syy019.

Carstens, B. C., S. J. Brunsfeld, J. R. Demboski, G. J. D, and J. Sullivan. 2005. Investigating the evolutionary history of the Pacific Northwest mesic forest ecosystem: hypothesis testing within a comparative phylogeographic framework. Evolution 59:1639–1652.

Castellanos, M. C., P. S. Wilson, S. J. Keller, A. D. Wolfe, and J. D. Thompson. 2006. Anther evolution: pollen presentation strategies when pollinators differ. American Naturalist 167:288–296.

Chifman, J. and L. S. Kubatko. 2015. Identifiability of the unrooted species tree topology under the coalescent model with time-reversible substitution processes, site-specific rate variation, and invariable sites. Journal of Theoretical Biology 374:35–47.

Crosswhite, F. S. 1965. Hybridization of *Penstemon barbatus* (Scrophulariaceae) of section *Elmigera* with species of *Habroanthus*. Southwestern Naturalist 10:234–237.

Datwyler, S. L. and A. D. Wolfe. 2004. Phylogenetic relationships and morphological evolution in *Penstemon* subg. Dasanthera (Veronicaceae). Systematic Botany 29:165–176.

Degnan, J. H. and N. A. Rosenberg. 2009. Gene tree discordance, phylogenetic inference and the multispecies coalescent. Trends in Ecology and Evolution 24:332–340.

Folk, R. A., J. R. Mandel, and J. V. Freudenstein. 2016. Ancestral gene flow and parallel organellar genome capture result in extreme phylogenomic discord in a lineage of angiosperms. Systematic Biology 66:320–337.

Folk, R. A., C. J. Visger, P. S. Soltis, D. E. Soltis, and R. P. Guralnick. 2018. Geographic range dynamics drove ancient hybridization in a lineage of angiosperms. American Naturalist https://dx.doi.org/10.1086/698120.

García, N., R. A. Folk, A. W. Meerow, S. Chamala, M. A. Gitzendanner, R. S. de Oliveira, D. E. Soltis, and P. S. Soltis. 2017. Deep reticulation and incomplete lineage sorting obscure the diploid phylogeny of rain-lillies and allies (Amaryllidaceae tribe Hippeastreae). Molecular Phylogenetics and Evolution 111:231–247.

Grant, V. 1971. Plant speciation. Columbia University Press.

Gruenstaeudl, M., N. M. Reid, G. L. Wheeler, and B. C. Carstens. 2015. Posterior predictive checks of coalescent models: P2C2M, an R package. Molecular Ecology Resources 16:193–205.

Jhwueng, D.-C. and B. O’Meara. 2015. Trait evolution on phylogenetic networks. bioRxiv https://doi.org/10.1101/023986.

Jones, G., S. Sagitov, and B. Oxelman. 2013. Statistical inference of allopolyploid species networks in the presence of incomplete lineage sorting. Systematic Biology 62:467–478.

Joshi, N. A. and J. N. Fash. 2011. Sickle: sliding-window, adaptive, quality-based trimming tool for FASTQ files (version 1.33). Available at https://github.com/najoshi/sickle.

Kamneva, O. K., J. Syring, A. Liston, and N. A. Rosenberg. 2017. Evaluating allopolyploid origins in strawberries (*Fragaria*) using haplotypes generated from target sequence capture. BMC Evolutionary Biology 17:180.

Katoh, S. 2013. MAFFT multiple sequence alignment software version 7: improvements in performance and usability. Molecular Biology and Evolution 30:772–780.

Keck, D. D. 1945. Studies in *Penstemon*–XIII: a cyto-taxonomic account of the section *Spermunculus*. American Midland Naturalist 33:128–206.

Kingman, J. F. C. 1982. On the genealogy of large populations. Journal of Applied Probability 19:27–43.

Kubatko, L. S. and J. Chifman. 2015. An invariants-based method for hybridization detection from genome-scale sequence data. bioRxiv https://doi.org/10.1101/034348.

Kubatko, L. S. and J. H. Degnan. 2007. Inconsistency of phylogenetic estimates from phylogenetic data under coalescence. Systematic Biology 56:17–24.

Larget, B., S. K. Kotha, C. N. Dewey, and C. Ané. 2010. BUCKy: gene tree / species tree reconciliation with Bayesian concordance analysis. Bioinformatics 26:2910–2911.

Lawrence, T. J. and S. L. Datwyler. 2016. Testing the hypothesis of allopolyploidy in the origin of *Penstemon azureus* (Plantaginaceae). Frontiers in Ecology and Evolution 4:60.

Levin, D. A. 1973. The role of trichomes in plant defense. Quarterly Review of Biology 48:3–15.

Lott, M., A. Spillner, K. T. Huber, and V. Moulton. 2009. PADRE: a package for analyzing and displaying reticulate evolution. Bioinformatics 25:1199–1200.

Maddison, W. P. 1997. Gene trees in species trees. Systematic Biology 46:523–536.

Magoč, T. and S. L. Salzberg. 2011. FLASH: fast length adjustment of short reads to improve genome assemblies. Bioinformatics 27:2957–2963.

Mallet, J. 2007. Hybrid speciation. Nature 446:279–283.

Marcussen, T., K. S. Jakobsen, J. Danihelka, H. E. Ballard, K. Blaxland, A. K. Brysting, and B. Oxelman. 2012. Inferring species networks from gene trees in high-polyploid North American and Hawaiian violets (*Viola*, Violaceae). Systematic Biology 61:107–126.

McDade, L. 1990. Hybrids and phylogenetic systematics I. Patterns of character expression in hybrids and their implications for cladistic analysis. Evolution 44:1685–1700.

McDade, L. 1992. Hybrids and phylogenetic systematics II. The impact of hybrids on cladistic analysis. Evolution 46:1329–1346.

Meng, C. and L. S. Kubatko. 2009. Detecting hybrid speciation in the presence of incomplete lineage sorting using gene tree incongruence: a model. Theoretical Population Biology 75:35–45.

Mirarab, S. and T. Warnow. 2015. ASTRAL-II: coalescent-based species tree estimation with many hundreds of taxa and thousands of genes. Bioinformatics 31:i44–i52.

Morales-Briones, D. F., A. Liston, and D. C. Tank. 2018. Phylogenomic analyses reveal a deep history of hybridization and polyploidy in the Neotropical genus *Lachemilla* (Rosaceae). New Phytologist https://doi.org/10.1111/nph.15099.

Nold, R. 1999. Penstemons. Timber Press, Portland, OR.

Oberpieler, C., W. F. S. Tomasello, and K. Konowalik. 2017. A permutation approach for inferring species networks from gene trees in polyploid complexes by minimizing deep coalescences. Methods in Ecology and Evolution 8:835–849.

Pamilo, P. and M. Nei. 1988. Relationships between gene trees and species trees. Molecular Biology and Evolution 5:568–583.

Reid, N. M., S. M. Hird, J. M. Brown, T. A. Pelletier, J. D. McVay, J. D. Satler, and B. C. Carstens. 2014. Poor fit to the multispecies coalescent is widely detectable in empirical data. Systematic Biology 63:322–333.

Rothfels, R. C., F.-W. Li, and K. M. Pryer. 2017. Next-generation polyploid phylogenetics: rapid resolution of hybrid polyploid complexes using PacBio single-molecule sequencing. New Phytologist 213:413–429.

Sayyari, E. and S. Mirarab. 2016. Fast coalescent-based computation of local branch support from quartet frequencies. Molecular Biology and Evolution 33:1654–1668.

Smith, S. A. and C. Dunn. 2008. Phyutility: a phyloinformatics utility for trees, alignments, and molecular data. Bioinformatics 24:715–716.

Snir, S. 2012. Quartet maxcut: a fast algorithm for amalgomating quartet trees. Molecular Phylogenetics and Evolution 62:1–8.

Solís-Lemus, C. and C. Ané. 2016. Inferring phylogenetic networks with maximum pseudolikelihood under incomplete lineage sorting. PLoS Genetics 12:e1005896.

Solís-Lemus, C., P. Bastide, and C. Ané. 2017. Phylonetworks: a package for phylogenetic networks. Molecular Biology and Evolution 34:3292–3298.

Stamatakis, A. 2014. RAxML version 8: a tool for phylogenetic analysis and post-analysis of large phylogenies. Bioinformatics 30:1312–1313.

Stamatakis, A., P. Hoover, and J. Rougemont. 2008. A rapid bootstrap algorithm for the RAxML web servers. Systematic Biology 57:758–771.

Stebbins, G. L. 1950. Variation and evolution in plants. Columbia University Press.

Stenz, N. W. M., B. Larget, D. A. Baum, and C. Ané. 2015. Exploring tree-like and non-tree-like patterns using genome sequences: an example using the inbreeding plant species *Arabisopsis thaliana* (L.) Heynh. Systematic Biology 64:809–823.

Straw, R. M. 1955. Hybridization, homogamy, and sympatric speciation. Evolution 9:441–444.

Straw, R. M. 1966. A redefinition of *Penstemon* (Scrophulariaceae). Brittonia 18:80–95.

Strickler, D. 1997. Northwest Penstemons. Flower Press.

Takahata, N. 1989. Gene genealogy in three related populations: consistency probability between gene and population trees. Genetics 122:957–966.

Tavaré, S. 1984. Line-of-descent and genealogical processes, and their applications in population genetics models. Theoretical Population Biology 26:119–164.

Uribe-Convers, S., M. L. Settles, and D. C. Tank. 2016. A phylogenomic approach based on PCR enrichment and high throughput sequencing: resolving diversity within the South American species of *Bartsia* L. (Orobanchaceae). PLoS ONE 11:e0148203.

Wen, D., Y. Yu, J. Zhu, and L. Nakhleh. 2018. Inferring phylogenetic networks using PhyloNet. Systematic Biology https://doi.org/10.1093/sysbio/syy015.

Wessinger, C. A., C. C. Freeman, M. E. Mort, M. D. Rausher, and L. C. Hileman. 2016. Multiplexed shotgun genotyping resolves species relationships within the North American genus *Penstemon*. American Journal of Botany 103:912–922.

Wilson, P. S., A. D. Wolfe, W. S. Armbruster, and J. D. Thompson. 2007. Constrained lability in floral evolution: counting convergent origins of hummingbird pollination in *Penstemon* and *Keckiella*. New Phytologist 176:883–890.

Wolfe, A. D. 2005. ISSR techniques for evolutionary biology. Methods in Enzymology 395:134–144.

Wolfe, A. D., S. L. Datwyler, and C. P. Randle. 2002. A phylogenetic and biogeographic analysis of the Cheloneae (Scrophulariaceae) based on ITS and matK sequence data. Systematic Botany 27:138–148.

Wolfe, A. D., C. P. Randle, S. L. Datwyler, J. J. Morawetz, N. Arguedas, and J. Diaz. 2006. Phylogeny, taxonomic affinities, and biogeography of *Penstemon* (Plantaginaceae) based on ITS and cpDNA sequence data. American Journal of Botany 93:1699–1713.

Wolfe, A. D., Q.-Y. Xiang, and S. R. Kephart. 1998a. Assessing hybridization in natural populations of *Penstemon* (Scrophulariaceae) using hypervariable intersimple sequence repeat (ISSR) bands. Molecular Ecology 7:1107–1125.

Wolfe, A. D., Q.-Y. Xiang, and S. R. Kephart. 1998b. Diploid hybrid speciation in *Penstemon* (Scrophulariaceae). Proceedings of the National Academy of Sciences 95:5112–5115.

