## Supplemental Materials for "Inferring Patterns of Hybridization and Polyploidy in the Plant Genus *Penstemon* (Plantaginaceae)"

Paul D. Blischak, Coleen E. Thompson, Emiko M. Waight,
Laura S. Kubatko, and Andrea D. Wolfe

### Supplemental Text

#### S1 Validating QCF Estimation

To evaluate the accuracy of our approach for QCF estimation, we performed a simulation study using both tree and network topologies (Figure 1), and compared our estimates with the true QCF values from simulated gene trees. Gene trees were simulated using the program `ms` for 50 loci using the specified topology with internal branch lengths of 0.5, 1.0, and 2.0 coalescent units (Hudson 2002). For the species network, the ancestral lineage to species C and D was simulated as a 60:40 hybrid species forming 1.0 coalescent units in the past through an admixture event between species E ($\gamma = 0.6$) and the ancestral lineage to A and B ($1 - \gamma = 0.4$). Sequence data was then simulated on each gene tree using the program Seq-Gen (Rambaut and Grass 1997). The length of each gene was 400 bp, with an expected number of substitutions per site of 0.05. QCFs were then estimated with the approach outlined in §2 (**Approach**) using either (1) no bootstrapping or (2) 500 bootstrap replicates. True QCF values were calculated using the simulated gene trees as input with the software package PhyloNetworks v0.7.0 (Solís-Lemus et al. 2017). Estimates from our approach were compared to the true values using the root mean squared deviation (RMSD), as well as linear regression, in R v3.3.2 (R Core Team 2016). Results were plotted using ggplot2 v2.2.1 (Wickham 2009). Code for performing these simulations can be found below in subsections §S1.1 and §S1.2.

##### S1.1 Tree Simulations

**qcf-sims-tree.sh:**

```
#!/bin/bash

# Global parameters
REP=$1                    # Rep number
THETA=0.05                # Expected number of mutations per base
BP=400                    # Sequence length in base pairs
julia5=/Applications/Julia-0.5.app/Contents/Resources/julia/bin/julia

for i in ‘seq 1 50‘
do
  # Simulate gene tree using ms
  ms 6 1 -T -I 6 1 1 1 1 1 1 -ej 0.25 1 2 -ej 0.25 3 4 -ej 0.5 2 4 \
     -ej 1.0 4 5 -ej 2.0 5 6 | grep ’^(’ > trees-${REP}-${i}.tre

  # Simulate sequence data using seq-gen
  seq-gen -mGTR -s $THETA -l $BP -r 1.0 0.2 10.0 0.75 3.2 1.6 \
     -f 0.15 0.35 0.15 0.35 -i 0.2 -a 5.0 -g 3 -q \
     < trees-${REP}-${i}.tre > seqs-${REP}-${i}.phy
done
```

```
ls -1 *.phy > genes.txt
cat trees-${REP}-*.tre > trees-${REP}.tre

qcf -i genes.txt -m map.txt --prefix tree-${REP}
qcf -i genes.txt -m map.txt -b 500 --prefix tree-${REP}-boot

$julia5 -e 'using PhyloNetworks; readTree2CF("trees-${REP}.tre", \
            "tree-${REP}-phynet.CFs.csv", writeSummary=false)'

mkdir rep-${REP}
mv *.tre *.phy *.csv rep-${REP}
```

```
$ for i in `seq 1 100`; do ./qcf-sims-tree.sh ${i}; done
```

#### S1.2 Network Simulations

**qcf-sims-network.sh:**

```
#!/bin/bash

# Global parameters
REP=$1                  # Rep number
THETA=0.05              # Expected number of mutations per base
BP=400                  # Sequence length in base pairs
julia5=/Applications/Julia-0.5.app/Contents/Resources/julia/bin/julia

# Simuate 20 gene trees from topology 1
for i in `seq 1 20`
do
  # Simulate gene tree using ms
  ms 6 1 -T -I 6 1 1 1 1 1 1 -ej 0.25 1 2 -ej 0.25 3 4 -ej 0.5 2 4 \
     -ej 1.0 4 5 -ej 2.0 5 6 | grep '^(' > trees-${REP}-${i}.tre

  # Simulate sequence data using seq-gen
  seq-gen -mGTR -s $THETA -l $BP -r 1.0 0.2 10.0 0.75 3.2 1.6 \
     -f 0.15 0.35 0.15 0.35 -i 0.2 -a 5.0 -g 3 -q \
     < trees-${REP}-${i}.tre > seqs-${REP}-${i}.phy
done

# Simulate 30 gene trees from topology 2
for i in `seq 21 50`
do
  # Simulate gene tree using ms
  ms 6 1 -T -I 6 1 1 1 1 1 1 -ej 0.25 1 2 -ej 0.25 3 4 -ej 0.5 4 5 \
     -ej 1.0 2 5 -ej 2.0 5 6 | grep '^(' > trees-${REP}-${i}.tre

  # Simulate sequence data using seq-gen
  seq-gen -mGTR -s $THETA -l $BP -r 1.0 0.2 10.0 0.75 3.2 1.6 \
     -f 0.15 0.35 0.15 0.35 -i 0.2 -a 5.0 -g 3 -q \
     < trees-${REP}-${i}.tre > seqs-${REP}-${i}.phy
done
```

```
ls *.phy > genes.txt
cat trees-${REP}-*.tre > trees-${REP}.tre

qcf -i genes.txt -m map.txt --prefix net-${REP}
qcf -i genes.txt -m map.txt -b 500 --prefix net-${REP}-boot

$julia5 -e 'using PhyloNetworks; readTree2CF("trees-${REP}.tre", \
            "net-${REP}-phynet.CFs.csv", writeSummary=false)'

mkdir rep-${REP}
mv *.tre *.phy *.csv rep-${REP}
```

```
$ for i in `seq 1 100`; do ./qcf-sims-network.sh ${i}; done
```

### S2 Code for Species Tree and Network Inference

#### S2.1 Gene Tree Estimates with RAxML

```
# Loop through all genes and analyze with RAxML
for f in *.phy
do
raxml -f a -x 12345 -p 12345 -# 500 -m GTRGAMMA \
      -s $f
done

# Then combine all gene trees
cat RAxML_bipartitions.* > AllGeneTrees.tre
```

#### S2.2 Species Tree Inference with ASTRAL-III

```
java -jar astral.5.5.9.jar -i AllGeneTrees.tre -a map.txt \
     -o Humiles-Proceri.tre --polylimit 20 \
     --samplingrounds 100 --extraLevel 2
```

The analyses for clades A and B were conducted using the same commands but with only the subset of taxa belonging to each clade.

##### S2.3 Species Tree Inference with `qcf`+QuartetMaxCut

```
# run QCF
qcf -i gene-list.txt -m map.txt -b 500

# Run get-pop-tree.pl from TICR
perl get-pop-tree.pl out-qcf.CFs.csv

# Run getTreeBranchLengths.R from TICR
Rscript getTreeBranchLengths.R out-qcf Pdavidsoniidavidsonii
```

##### S2.4 Network Analyses with PhyloNetworks

Network analyses were conducted using the SNaQ method in the PhyloNetworks package (v0.7.0) with the following template for each script [written in the Julia language using versions 5.2.0 and 6.2.0] (Solís-Lemus and Ané 2016; Solís-Lemus et al. 2017).

```
# snaq-net<H>.jl
addprocs(10) # add processors to run things in parallel
using Phylonetworks;

t  = readTopology("cladeA-astral.tre")
cf = readTableCF("cladeA-qcf.CFs.csv")
net1 = snaq!(t, cf, hmax=<H>,
             filename="cladA-net<H>",
             outgroup="Pdavidsoniidavidsonii")
```

For each network analysis, we changed the `hmax` argument to the corresponding maximum number of hybridization events (`<H>` = 1 through 5).

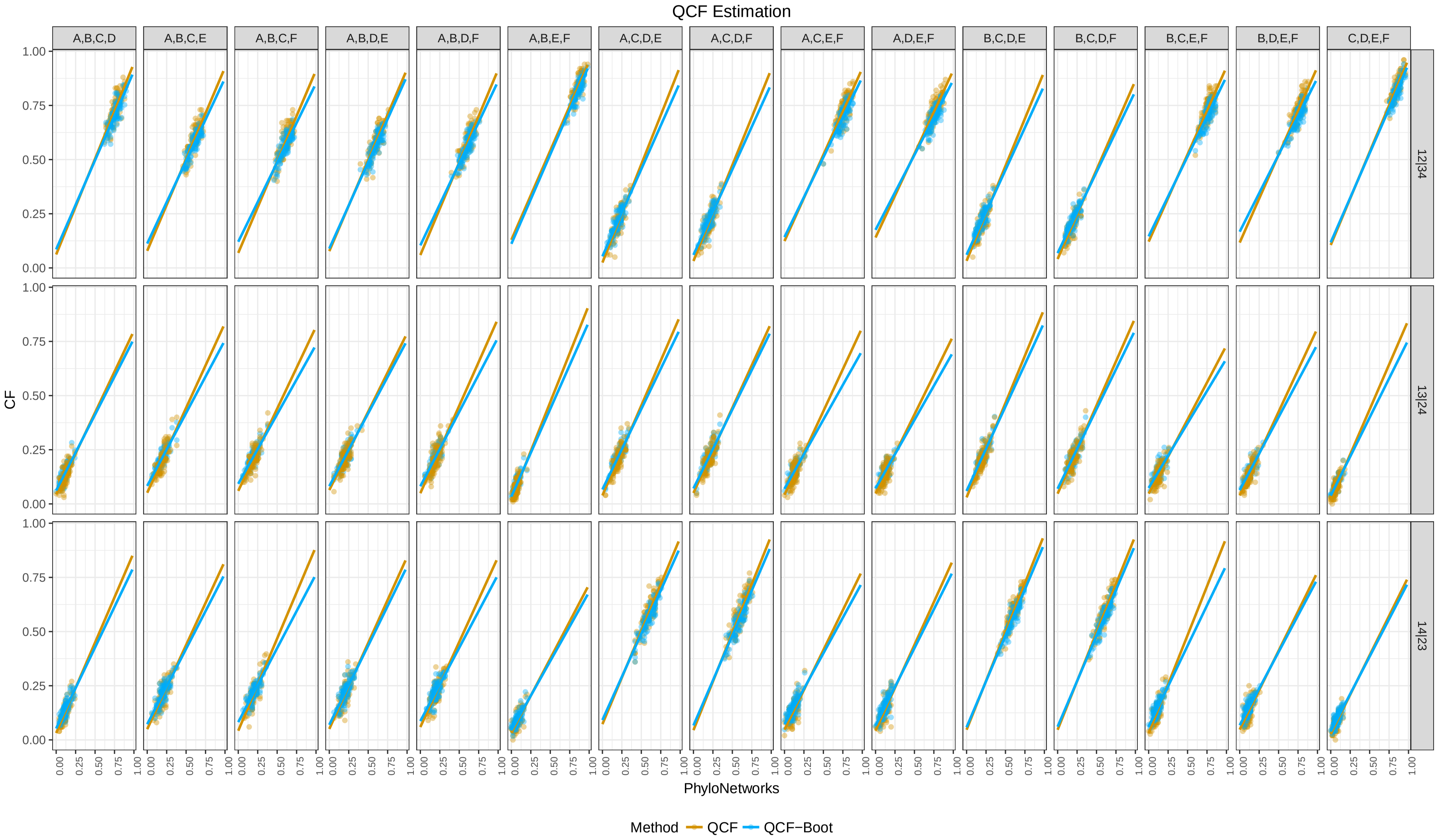

**Fig. S1.** Simulation results for the tree topology from Figure 1a. Results are plotted for bootstrapped (blue) and non-bootstrapped (yellow) comparisons. Regression lines were estimated by ggplot2 using the `lm` method in the `geom_smooth()` function.

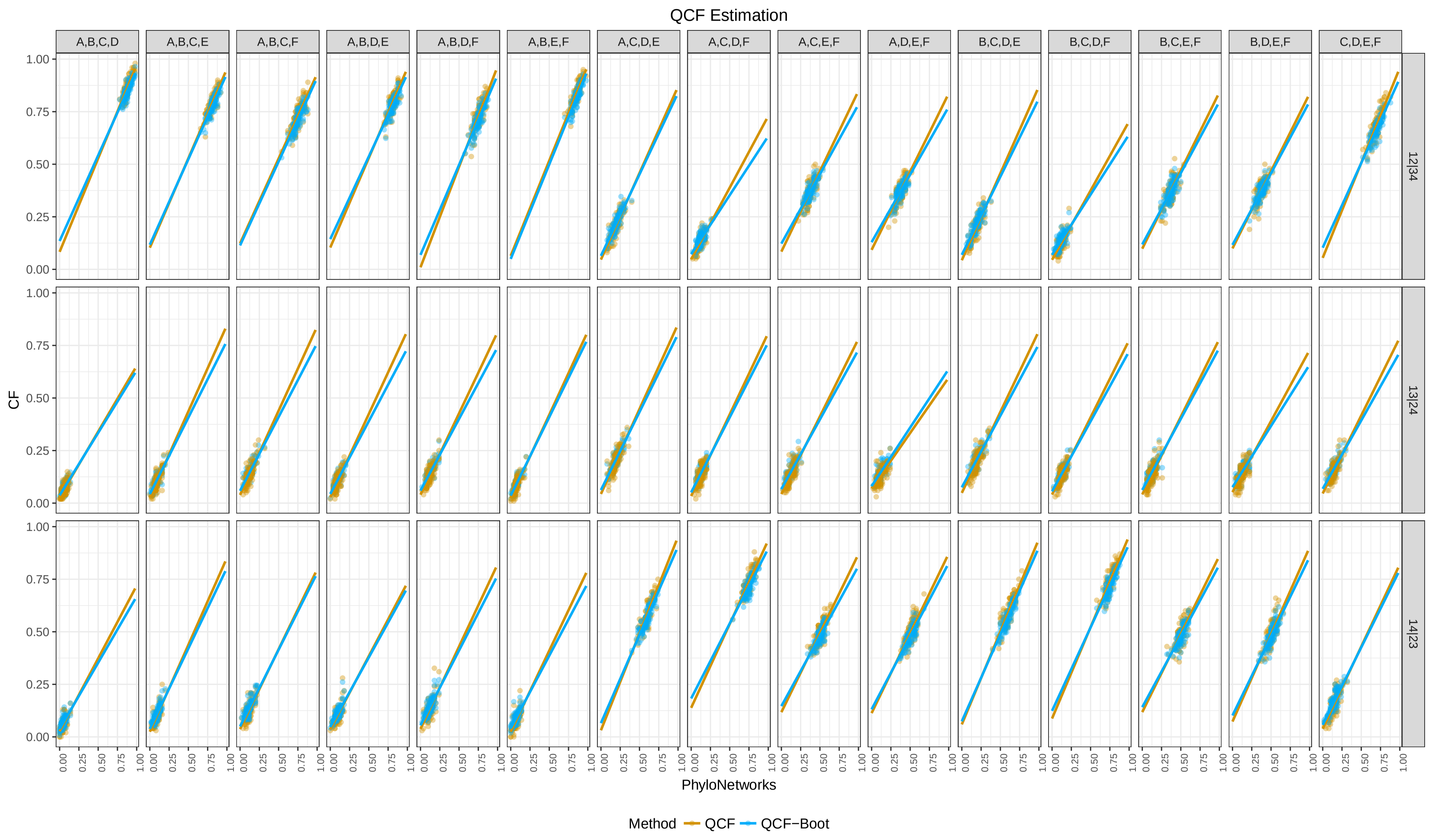

**Fig. S2.** Simulation results for the network topology from Figure 1b. Results are plotted for bootstrapped (blue) and non-bootstrapped (yellow) comparisons. Regression lines were estimated by ggplot2 using the `lm` method in the `geom_smooth()` function.

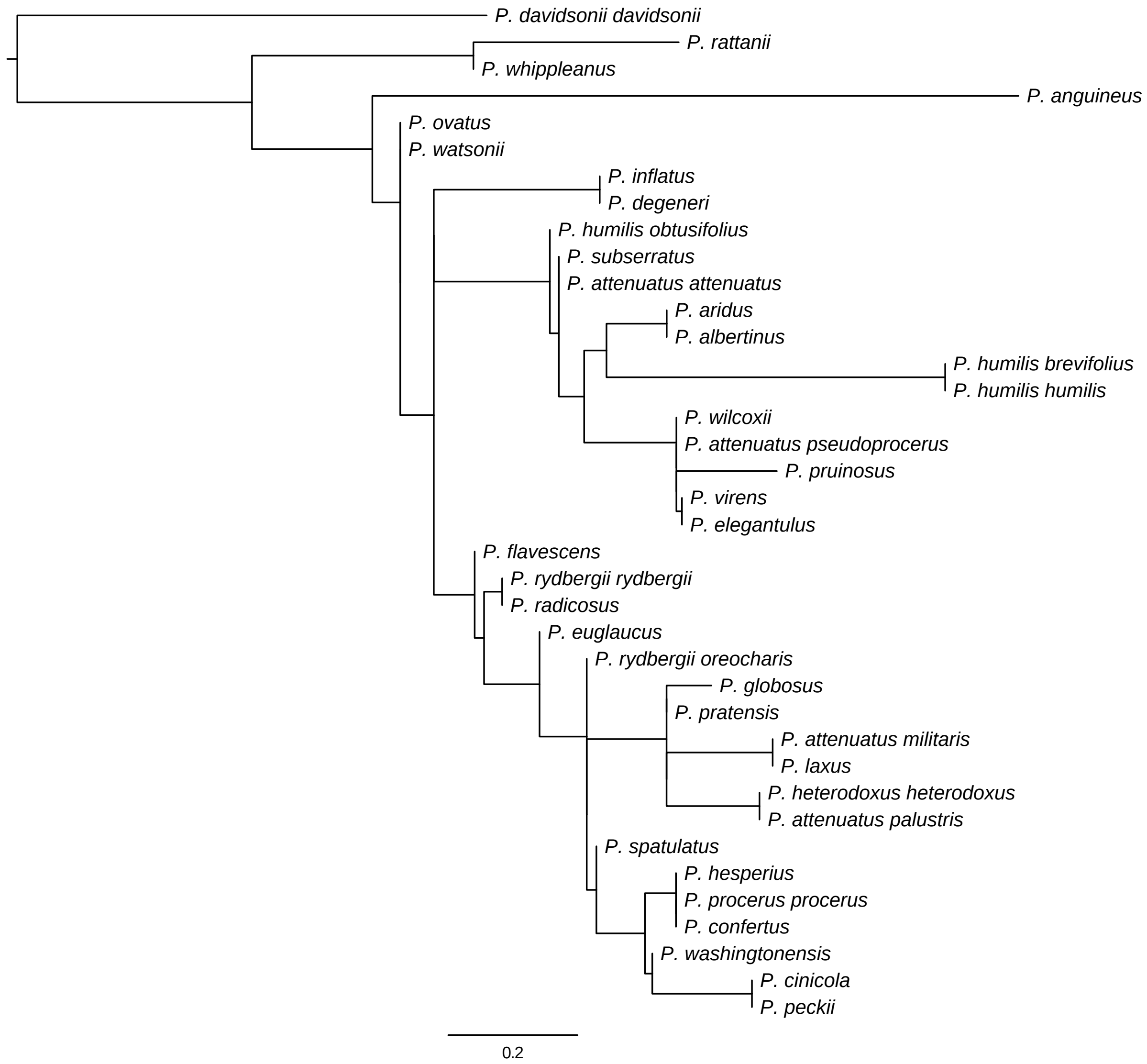

**Fig. S3.** Phylogeny of *Penstemon* subsections *Humiles* and *Proceri* inferred by ASTRAL-III. Branch lengths are in coalescent units.

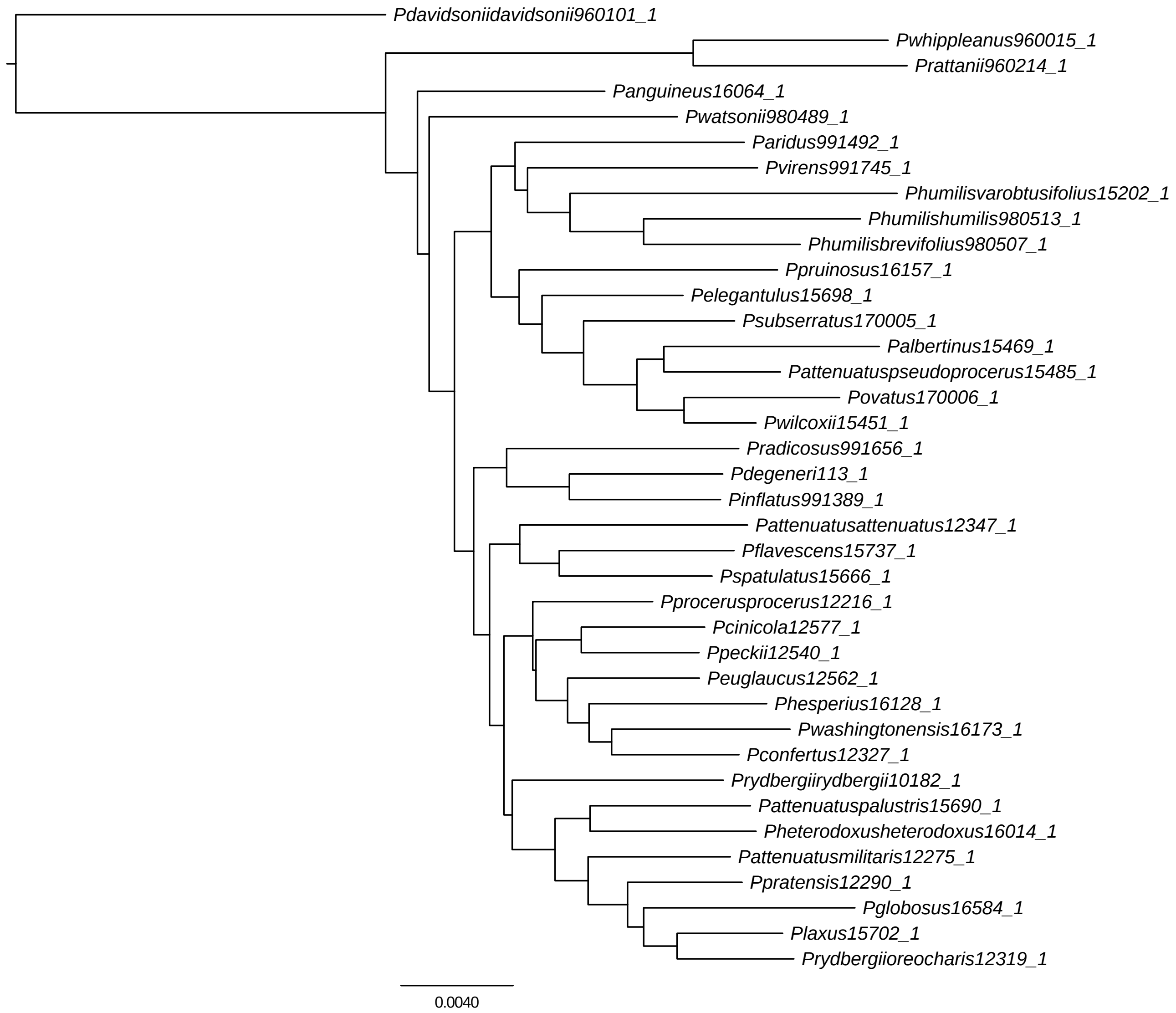

**Fig. S4.** Phylogeny of *Penstemon* subsections *Humiles* and *Proceri* inferred with RAxML v8.2.11 using a supermatrix of 43 loci. Branch lengths are the mean number of substitutions per site.

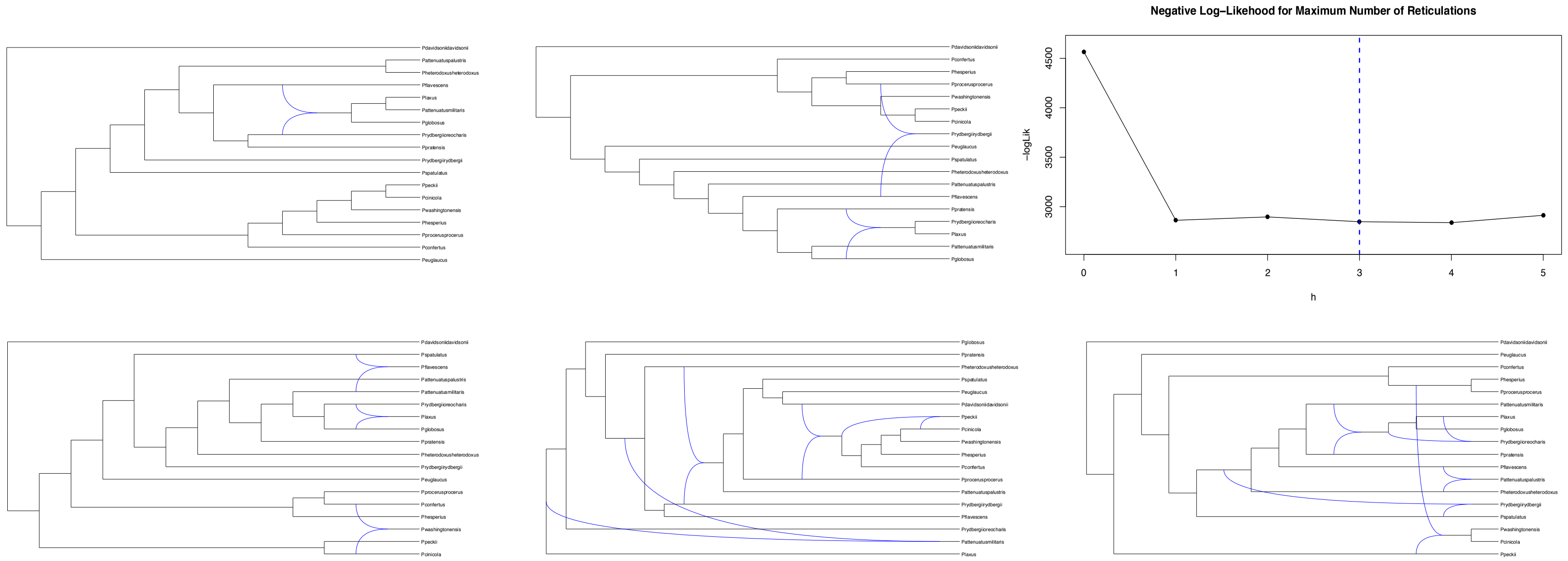

**Fig. S5.** Networks inferred for clade A using SNaQ, as implemented in the software PhyloNetworks. Each panel shows the network estimated for the given number of hybridization events (h=1 to h=5), starting at the top left (h=1) and ending at the bottom right (h=5). The panel in the top right corner shows the negative log-pseduolikelihoods for each number of hybridization events, with h=3 having the best pseduolikelihood.

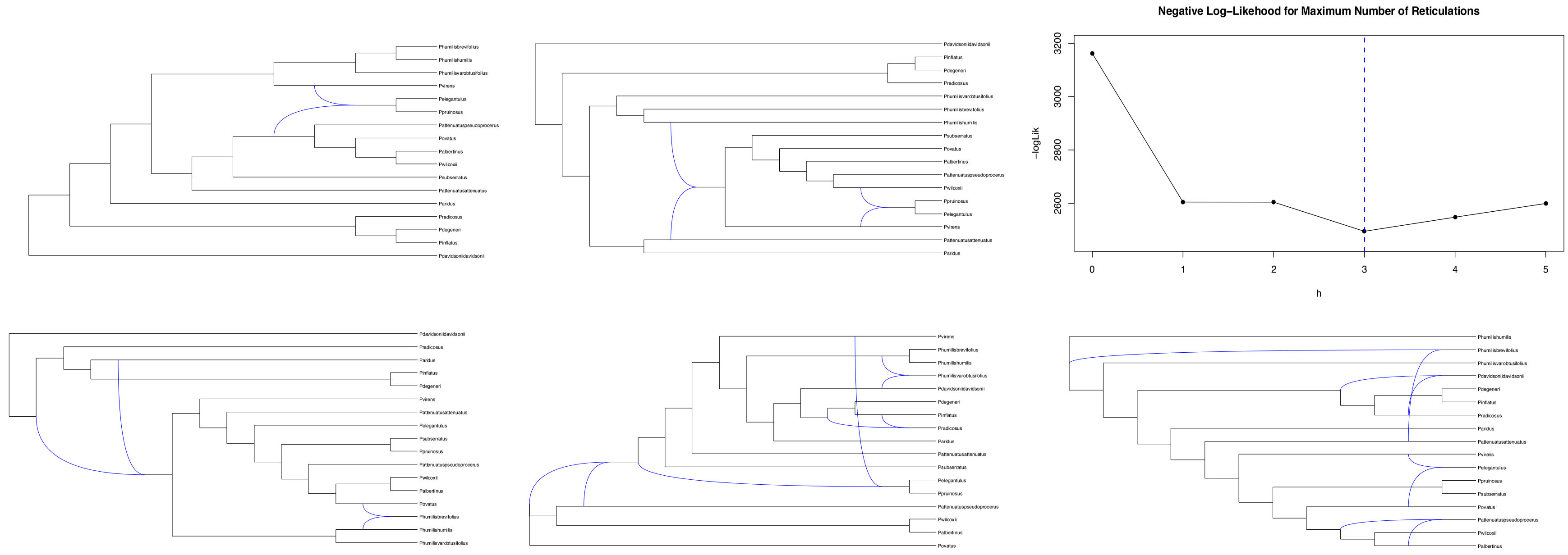

**Fig. S6.** Networks inferred for clade B using SNaQ, as implemented in the software PhyloNetworks. Each panel shows the network estimated for the given number of hybridization events (h=1 to h=5), starting at the top left (h=1) and ending at the bottom right (h=5). The panel in the top right corner shows the negative log-pseduolikelihoods for each number of hybridization events, with h=3 having the best pseduolikelihood.

**Table S1.** Collection and ploidy information for accessions from *Penstemon* subsections *Humiles* and *Proceri*.

| Taxon | Ploidy | Voucher |
|---|---|---|
| ***Outgroup*** | | |
| *P. davidsonii davidsonii* | 2X | SLD 53 |
| ***Subsection* Humiles** | | |
| *P. albertinus* | 2X | PDB 41 |
| *P. anguineus* | 2X | ADW 1505 |
| *P. aridus* | 2X | Andrew Lutz, Billy Creek 1 |
| *P. degeneri* | 2X | ADW 401 |
| *P. elegantulus* | 2X | Idaho Gray 4321 |
| *P. inflatus* | 2X | ADW 811 |
| *P. humilis brevifolius* | 2X | ADW 573 |
| *P. humilis humilis* | 2X | ADW 761 |
| *P. humilis obtusifolius* | 2X | ADW 1430 |
| *P. ovatus* | 2X | ADW 608 |
| *P. pruinosus* | 2X | BYU 98608/EC Moran sn (1971) |
| *P. radicosus* | 2X | Mt. West Enviro. Services 7962 |
| *P. rattanii* | 2X | ADW 512 |
| *P. subserratus* | 2X | ADW 590 |
| *P. virens* | 2X | Mt. West Enviro. Services 7953 |
| *P. whippleanus* | 2X | 1073 Potsdam |
| *P. wilcoxii* | 2X | PDB 39 |
| ***Subsection* Proceri** | | |
| *P. attenuatus attenuatus* | 6X | PDB 19 |
| *P. attenuatus militaris* | 6X | PDB 12 |
| *P. attenuatus palustris* | 6X | PDB 61 |
| *P. attenuatus pseudoprocerus* | 6X | PDB 42 |
| *P. cinicola* | 2X | PDB 32 |
| *P. confertus* | 4X | PDB 18 |
| *P. euglaucus* | 6X | PDB 33 |
| *P. flavescens* | 6X | PDB 63 |
| *P. globosus* | 4X | ADW 1566 |
| *P. hesperius* | 2X | – |
| *P. heterodoxus heterodoxus* | 2X | ADW 1498 |
| *P. laxus* | 2X | Idaho Smith 8123 |
| *P. peckii* | 4X | PDB 31 |
| *P. pratensis* | 2X | PDB 14 |
| *P. procerus procerus* | 4X | PDB 3 |
| *P. rydbergii rydbergii* | 4X | Western Env. 8845 |
| *P. rydbergii oreocharis* | 2X | PDB 16 |
| *P. spatulatus* | 2X | PDB 56 |
| *P. washingtonensis* | 2X | PDB 36 |
| *P. watsonii* | 2X | ADW 786 |

**Table S2.** Primers for amplicon sequencing using using the Fluidigm AccessArray. Primer sequences include conserved sequence tags.

| Locus | Direction | Primer |
|---|---|---|
| COS4270 | Forward | ACACTGACGACATGGTTCTACAACCAAGCTCTTCACCTGGAA |
| | Reverse | TACGGTAGCAGAGACTTGGTCTAGACCAGCATAACAATTTTATTCCTAA |
| COS14240 | Forward | ACACTGACGACATGGTTCTACACCGACATTAGTCACGGTCCT |
| | Reverse | TACGGTAGCAGAGACTTGGTCTCGGCATTCCTTCAGATAAAC |
| COS23460 | Forward | ACACTGACGACATGGTTCTACATGGTTGTTCGTGTGAGGTTG |
| | Reverse | TACGGTAGCAGAGACTTGGTCTAAACTCATTATTTGCTCGATAGGG |
| COS24530 | Forward | ACACTGACGACATGGTTCTACATTAAAATGCAGGAGGGCTTG |
| | Reverse | TACGGTAGCAGAGACTTGGTCTCCAACCAAATCAGTTCTCTGC |
| COS50360 | Forward | ACACTGACGACATGGTTCTACACCATGGAATCAAACCTGGAC |
| | Reverse | TACGGTAGCAGAGACTTGGTCTAAGCCCAAATCGAAGAAGAA |
| COS57850 | Forward | ACACTGACGACATGGTTCTACAGAAGGAGCCTCAAAGCAGTG |
| | Reverse | TACGGTAGCAGAGACTTGGTCTGGATGTCCATCTAACCCGTTT |
| PPR876 | Forward | ACACTGACGACATGGTTCTACACAGCTTCTGGTAGATGGGCT |
| | Reverse | TACGGTAGCAGAGACTTGGTCTCTCTCCCCACAATCTTCGCC |
| PPR1651 | Forward | ACACTGACGACATGGTTCTACAACGCAGCTTCTGGTAGATGG |
| | Reverse | TACGGTAGCAGAGACTTGGTCTCACCACAATTACACACGCCC |
| PPR5729 | Forward | ACACTGACGACATGGTTCTACATTTGCTCACTGCTTGTGCTG |
| | Reverse | TACGGTAGCAGAGACTTGGTCTCCTTCATCCACCATGCCACA |
| PPR985 | Forward | ACACTGACGACATGGTTCTACATCCGTGGCATTTTTAGTGCG |
| | Reverse | TACGGTAGCAGAGACTTGGTCTCCGGAGAAAGCTCTTACGGTT |
| PPR1839 | Forward | ACACTGACGACATGGTTCTACATGCAACCGTAATGCTCGACT |
| | Reverse | TACGGTAGCAGAGACTTGGTCTTTTGCTCACTGCTTGTGCTG |
| 34130 | Forward | ACACTGACGACATGGTTCTACATCTAAGTTTGCGGATGTTGAGA |
| | Reserve | TACGGTAGCAGAGACTTGGTCTCATTCCCAGAACATACATGCAA |
| 59820 | Forward | ACACTGACGACATGGTTCTACAGCAGATTTAGTTTTACTCTCCTCCA |
| | Reverse | TACGGTAGCAGAGACTTGGTCTGGTCTTAAATACCATCTTCTGTGTCC |
| 80460 | Forward | ACACTGACGACATGGTTCTACACCGAAATTTTACCCAAAATCG |
| | Reverse | TACGGTAGCAGAGACTTGGTCTGCAATGTGGGATTTGTTCGT |
| 20370 | Forward | ACACTGACGACATGGTTCTACATTCAGAGCTCCCATTTTTGC |
| | Reverse | TACGGTAGCAGAGACTTGGTCTTTGACCTTCATCCAATAGAGCA |
| 21370 | Forward | ACACTGACGACATGGTTCTACACTGTTTTTCCAATTTTCCATCC |
| | Reverse | TACGGTAGCAGAGACTTGGTCTCAGGTTGTGGGCTACGATTT |
| PPR369 | Forward | ACACTGACGACATGGTTCTACAGGAAAGGAAATCCATGCCCA |
| | Reverse | TACGGTAGCAGAGACTTGGTCTAGCCTTCAGTTACCATTCCG |
| PPR950 | Forward | ACACTGACGACATGGTTCTACATCTCCATCTTCGAGAACGCC |
| | Reverse | TACGGTAGCAGAGACTTGGTCTGCCGGATCAGGTGACGATAG |
| PPR1250 | Forward | ACACTGACGACATGGTTCTACAAAAGCCCTTCTTGCACGAGT |
| | Reverse | TACGGTAGCAGAGACTTGGTCTGGACAGCTTTGATTGCAGGG |
| PPR1561 | Forward | ACACTGACGACATGGTTCTACATCCCTTTTGCCTCATCGACC |
| | Reverse | TACGGTAGCAGAGACTTGGTCTGGTTCACACGGTGAATGTCG |
| 30360 | Forward | ACACTGACGACATGGTTCTACAAGGTTGCTAAAGGCCGATTC |
| | Reverse | TACGGTAGCAGAGACTTGGTCTGGGTCTTTATCTAAAAGGCGAGA |
| 35920 | Forward | ACACTGACGACATGGTTCTACAGGGGACAAAAATAGCAGAGC |
| | Reverse | TACGGTAGCAGAGACTTGGTCTTCACCGTGCTTGTTAAGTGC |
| 27260 | Forward | ACACTGACGACATGGTTCTACACTCCCCCGGAAAGTAACAAA |
| | Reverse | TACGGTAGCAGAGACTTGGTCTTTGTTTCATGTTGCGCCTTT |
| 2840 | Forward | ACACTGACGACATGGTTCTACATCTGGAAAAATTCCCTGGAC |
| | Reverse | TACGGTAGCAGAGACTTGGTCTGCGCTTTGCAAATTCTTGAG |

**Table S3.** Primers for amplicon sequencing using using the Fluidigm AccessArray (continued). Primer sequences include conserved sequence tags.

| Locus | Direction | Primer |
|---|---|---|
| 53950 | Forward | ACACTGACGACATGGTTCTACAAAACTGTGCTCTTCCTCCAA |
| | Reverse | TACGGTAGCAGAGACTTGGTCTGGGGAATGGTGACTCCTACA |
| 62010 | Forward | ACACTGACGACATGGTTCTACAAGCAACGCCATAAACTGGAA |
| | Reverse | TACGGTAGCAGAGACTTGGTCTTTGGGAAAGTTGATTGAGACG |
| 2350 | Forward | ACACTGACGACATGGTTCTACAGCTCCCATCTTTGTATATCTCG |
| | Reverse | TACGGTAGCAGAGACTTGGTCTCCCGTTCGTTCGATTGATAG |
| 18520 | Forward | ACACTGACGACATGGTTCTACATTCGAGAACGCCTCTAAACC |
| | Reverse | TACGGTAGCAGAGACTTGGTCTGTTATGCACAAAACGGGATG |
| 33495 | Forward | ACACTGACGACATGGTTCTACATCTCCATTTCTCAACCTCAGC |
| | Reverse | TACGGTAGCAGAGACTTGGTCTGCCCCTTCCTCTCCATACA |
| 38180 | Forward | ACACTGACGACATGGTTCTACAGGCATCAAAAGTGGATGATG |
| | Reverse | TACGGTAGCAGAGACTTGGTCTTCCCTCGTTGAGACATTCCT |
| 4810 | Forward | ACACTGACGACATGGTTCTACATACCAATTCCCCAGTTCTGC |
| | Reverse | TACGGTAGCAGAGACTTGGTCTATGGGAAGAGATGCTTACCTGA |
| 37450 | Forward | ACACTGACGACATGGTTCTACATGCTATCAAAACTTCGGCATC |
| | Reverse | TACGGTAGCAGAGACTTGGTCTGATCTCAAAAAGCACAACTCCA |
| 66330 | Forward | ACACTGACGACATGGTTCTACATGCAAATTCTTGAGCTGTCC |
| | Reverse | TACGGTAGCAGAGACTTGGTCTAAAATTCCCTGGACGCTTG |
| 1331 | Forward | ACACTGACGACATGGTTCTACAGCACCAAGATATGCCATTGA |
| | Reverse | TACGGTAGCAGAGACTTGGTCTTCCGAGCTAAGGCTATACATTCA |
| 48730 | Forward | ACACTGACGACATGGTTCTACACCATACGCGTAAATAAGAGAGC |
| | Reverse | TACGGTAGCAGAGACTTGGTCTTGATGGATATGGTAAAGCTAAACG |
| 3382 | Forward | ACACTGACGACATGGTTCTACATCTGAAAGCCTTGTACCAACC |
| | Reverse | TACGGTAGCAGAGACTTGGTCTGAGCCCTCTTGCCATTTCTA |
| 2829 | Forward | ACACTGACGACATGGTTCTACATGACCCGTTGACAACCCTAT |
| | Reverse | TACGGTAGCAGAGACTTGGTCTTGCTTACAGGCCCTTTGGTA |
| 604 | Forward | ACACTGACGACATGGTTCTACAATGGCCTCCGGTAATTCTCT |
| | Reverse | TACGGTAGCAGAGACTTGGTCTAGTGGCCTGAACTTTGCAGT |
| 598 | Forward | ACACTGACGACATGGTTCTACATCTGGGCTAACCTGAAATCG |
| | Reverse | TACGGTAGCAGAGACTTGGTCTTGGAAATGATCAAGAAATGAAGC |
| 233 | Forward | ACACTGACGACATGGTTCTACAACAACGCTGTGTGTTTGGTC |
| | Reverse | TACGGTAGCAGAGACTTGGTCTCCCACCAGCCCTTAACTACTC |
| 2978 | Forward | ACACTGACGACATGGTTCTACATCCATACAATGGTAAGATCACAAGA |
| | Reverse | TACGGTAGCAGAGACTTGGTCTGAAGGGTGTTCCGGGATTAT |
| 2919 | Forward | ACACTGACGACATGGTTCTACAAGAGTGTCATGGCCACCAAT |
| | Reverse | TACGGTAGCAGAGACTTGGTCTATGGACCGCATAGCTCAAAG |
| 2782 | Forward | ACACTGACGACATGGTTCTACAGAATTGAGGAGATTTGGGAATTT |
| | Reverse | TACGGTAGCAGAGACTTGGTCTCAGAATTGGGCCCTCCTAAG |
| 1397 | Forward | ACACTGACGACATGGTTCTACATCCAGTTTCGCTGAAATCACT |
| | Reverse | TACGGTAGCAGAGACTTGGTCTTAAAGGCCTTGGAGAAGCAA |
| 836 | Forward | ACACTGACGACATGGTTCTACATCCCAATTTATCCCAGAAAGC |
| | Reverse | TACGGTAGCAGAGACTTGGTCTAATCATGGGCGACCTATTTG |
| rps12rpl20 | Forward | ACACTGACGACATGGTTCTACAATTAGAAANRCAAGACAGCCAAT |
| | Reverse | TACGGTAGCAGAGACTTGGTCTCGYYAYCGAGCTATATATCC |
| trnTL | Forward | ACACTGACGACATGGTTCTACACATTACAAATGCGATGCTCT |
| | Reverse | TACGGTAGCAGAGACTTGGTCTTCTACCGATTTCGCCATATC |
| trnCD | Forward | ACACTGACGACATGGTTCTACACCAGTTCAAATCTGGGTGTC |
| | Reverse | TACGGTAGCAGAGACTTGGTCTGGGATTGTAGTTCAATTGGT |

**Table S4.** RMSD values for QCF estimation using data simulated from a tree topology (Figure 1a).

| Quartet | Topology | QCF | QCF-Boot |
|---|---|---|---|
| A,B,C,D | 12 \| 34 | 0.0365333 | 0.0310762 |
| A,B,C,D | 13 \| 24 | 0.0283334 | 0.0293331 |
| A,B,C,D | 14 \| 23 | 0.0242000 | 0.0524701 |
| A,B,C,E | 12 \| 34 | 0.0336667 | 0.0307411 |
| A,B,C,E | 13 \| 24 | 0.0295333 | 0.0279207 |
| A,B,C,E | 14 \| 23 | 0.0304000 | 0.0357291 |
| A,B,C,F | 12 \| 34 | 0.0423667 | 0.0334601 |
| A,B,C,F | 13 \| 24 | 0.0314000 | 0.0293172 |
| A,B,C,F | 14 \| 23 | 0.0297667 | 0.0465771 |
| A,B,D,E | 12 \| 34 | 0.0320666 | 0.0301693 |
| A,B,D,E | 13 \| 24 | 0.0313667 | 0.0285252 |
| A,B,D,E | 14 \| 23 | 0.0311000 | 0.0377345 |
| A,B,D,F | 12 \| 34 | 0.0387000 | 0.0317376 |
| A,B,D,F | 13 \| 24 | 0.0317334 | 0.0317221 |
| A,B,D,F | 14 \| 23 | 0.0319667 | 0.0447368 |
| A,B,E,F | 12 \| 34 | 0.0307667 | 0.0247832 |
| A,B,E,F | 13 \| 24 | 0.0233333 | 0.0252398 |
| A,B,E,F | 14 \| 23 | 0.0229000 | 0.0394890 |
| A,C,D,E | 12 \| 34 | 0.0282000 | 0.0265650 |
| A,C,D,E | 13 \| 24 | 0.0282333 | 0.0365885 |
| A,C,D,E | 14 \| 23 | 0.0323000 | 0.0257895 |
| A,C,D,F | 12 \| 34 | 0.0306667 | 0.0286579 |
| A,C,D,F | 13 \| 24 | 0.0314666 | 0.0411249 |
| A,C,D,F | 14 \| 23 | 0.0361333 | 0.0283903 |
| A,C,E,F | 12 \| 34 | 0.0387333 | 0.0313787 |
| A,C,E,F | 13 \| 24 | 0.0293333 | 0.0382125 |
| A,C,E,F | 14 \| 23 | 0.0329333 | 0.0578481 |
| A,D,E,F | 12 \| 34 | 0.0399333 | 0.0323120 |
| A,D,E,F | 13 \| 24 | 0.0290667 | 0.0377330 |
| A,D,E,F | 14 \| 23 | 0.0312000 | 0.0587910 |
| B,C,D,E | 12 \| 34 | 0.0268000 | 0.0258273 |
| B,C,D,E | 13 \| 24 | 0.0245333 | 0.0355890 |
| B,C,D,E | 14 \| 23 | 0.0260000 | 0.0280845 |
| B,C,D,F | 12 \| 34 | 0.0247333 | 0.0312867 |
| B,C,D,F | 13 \| 24 | 0.0313000 | 0.0387545 |
| B,C,D,F | 14 \| 23 | 0.0295667 | 0.0284660 |
| B,C,E,F | 12 \| 34 | 0.0365333 | 0.0315087 |
| B,C,E,F | 13 \| 24 | 0.0272000 | 0.0346498 |
| B,C,E,F | 14 \| 23 | 0.0296667 | 0.0532958 |
| B,D,E,F | 12 \| 34 | 0.0360000 | 0.0285520 |
| B,D,E,F | 13 \| 24 | 0.0244000 | 0.0334152 |
| B,D,E,F | 14 \| 23 | 0.0313333 | 0.0534680 |
| C,D,E,F | 12 \| 34 | 0.0252000 | 0.0228099 |
| C,D,E,F | 13 \| 24 | 0.0194000 | 0.0227021 |
| C,D,E,F | 14 \| 23 | 0.0207333 | 0.0345828 |

**Table S5.** RMSD values for QCF estimation using data simulated from a network topology (Figure 1b).

| Quartet | Topology | QCF | QCF-Boot |
|---|---|---|---|
| A,B,C,D | 12 \| 34 | 0.0232667 | 0.0213276 |
| A,B,C,D | 13 \| 24 | 0.0199333 | 0.0212925 |
| A,B,C,D | 14 \| 23 | 0.0191333 | 0.0323620 |
| A,B,C,E | 12 \| 34 | 0.0255000 | 0.0235640 |
| A,B,C,E | 13 \| 24 | 0.0211000 | 0.0240413 |
| A,B,C,E | 14 \| 23 | 0.0219333 | 0.0345390 |
| A,B,C,F | 12 \| 34 | 0.0342000 | 0.0277485 |
| A,B,C,F | 13 \| 24 | 0.0274000 | 0.0242728 |
| A,B,C,F | 14 \| 23 | 0.0261333 | 0.0412771 |
| A,B,D,E | 12 \| 34 | 0.0277000 | 0.0237231 |
| A,B,D,E | 13 \| 24 | 0.0225000 | 0.0239117 |
| A,B,D,E | 14 \| 23 | 0.0235333 | 0.0328303 |
| A,B,D,F | 12 \| 34 | 0.0341000 | 0.0275928 |
| A,B,D,F | 13 \| 24 | 0.0246667 | 0.0279635 |
| A,B,D,F | 14 \| 23 | 0.0277000 | 0.0425720 |
| A,B,E,F | 12 \| 34 | 0.0245000 | 0.0222437 |
| A,B,E,F | 13 \| 24 | 0.0190000 | 0.0241789 |
| A,B,E,F | 14 \| 23 | 0.0205000 | 0.0381137 |
| A,C,D,E | 12 \| 34 | 0.0266000 | 0.0233988 |
| A,C,D,E | 13 \| 24 | 0.0270334 | 0.0358624 |
| A,C,D,E | 14 \| 23 | 0.0281000 | 0.0273872 |
| A,C,D,F | 12 \| 34 | 0.0243667 | 0.0259449 |
| A,C,D,F | 13 \| 24 | 0.0287000 | 0.0369820 |
| A,C,D,F | 14 \| 23 | 0.0288667 | 0.0292340 |
| A,C,E,F | 12 \| 34 | 0.0294000 | 0.0330291 |
| A,C,E,F | 13 \| 24 | 0.0273000 | 0.0286997 |
| A,C,E,F | 14 \| 23 | 0.0281667 | 0.0265154 |
| A,D,E,F | 12 \| 34 | 0.0311333 | 0.0349219 |
| A,D,E,F | 13 \| 24 | 0.0323667 | 0.0313181 |
| A,D,E,F | 14 \| 23 | 0.0313000 | 0.0244409 |
| B,C,D,E | 12 \| 34 | 0.0267667 | 0.0263707 |
| B,C,D,E | 13 \| 24 | 0.0282000 | 0.0345829 |
| B,C,D,E | 14 \| 23 | 0.0311667 | 0.0280294 |
| B,C,D,F | 12 \| 34 | 0.0264667 | 0.0263840 |
| B,C,D,F | 13 \| 24 | 0.0264000 | 0.0366836 |
| B,C,D,F | 14 \| 23 | 0.0268000 | 0.0291028 |
| B,C,E,F | 12 \| 34 | 0.0318667 | 0.0311415 |
| B,C,E,F | 13 \| 24 | 0.0287333 | 0.0305662 |
| B,C,E,F | 14 \| 23 | 0.0301333 | 0.0293290 |
| B,D,E,F | 12 \| 34 | 0.0300333 | 0.0340250 |
| B,D,E,F | 13 \| 24 | 0.0313667 | 0.0315823 |
| B,D,E,F | 14 \| 23 | 0.0358000 | 0.0246551 |
| C,D,E,F | 12 \| 34 | 0.0295333 | 0.0258286 |
| C,D,E,F | 13 \| 24 | 0.0241667 | 0.0287898 |
| C,D,E,F | 14 \| 23 | 0.0299666 | 0.0373072 |

#### Supplemental References

Hudson, R. R. 2002. Generating samples under a Wright-Fisher neutral model of genetic variation. Bioinformatics 18:337–338.

R Core Team. 2016. R: a language and environment for statistical computing. R Foundation for Statistical Computing Vienna, Austria.

Rambaut, A. and N. C. Grass. 1997. Seq-gen: an application for the monte carlo simulation of dna sequence evolution along phylogenetic trees. Computer Applications in the Biosciences 13:235–238.

Solís-Lemus, C. and C. Ané. 2016. Inferring phylogenetic networks with maximum pseudolikelihood under incomplete lineage sorting. PLoS Genetics 12:e1005896.

Solís-Lemus, C., P. Bastide, and C. Ané. 2017. Phylonetworks: a package for phylogenetic networks. Molecular Biology and Evolution 34:3292–3298.

Wickham, H. 2009. ggplot2: elegant graphics for data analysis. Springer, New York.
